# Erosion of X-Chromosome Inactivation in female hiPSCs is heterogeneous and persists during differentiation

**DOI:** 10.1101/2024.03.15.585169

**Authors:** Ana Cláudia Raposo, Paulo Caldas, Maria Arez, Joana Jeremias, Pedro Barbosa, Rui Sousa-Luís, Frederico Água, David Oxley, Annalisa Mupo, Melanie Eckersley-Maslin, Miguel Casanova, Ana Rita Grosso, Simão Teixeira da Rocha

## Abstract

During culture, female human pluripotent stem cells (hPSCs), including human induced PSCs (hiPSCs) exhibit a propensity for erosion of X-chromosome inactivation (XCI). This phenomenon is characterized by the loss of XIST RNA expression and reactivation of a subset of X-linked genes from the inactive X chromosome (Xi). XCI erosion, despite its common occurrence, is often overlooked by the stem cell community, hindering a complete understanding of its impact on both fundamental and translational applications of hiPSCs. Investigating erosion dynamics in female hiPSCs, our study reveals that XCI erosion is a frequent yet heterogeneous phenomenon, resulting in the reactivation of several X-linked genes. The likelihood of a gene to erode increases for those located on the short arm of the X chromosome and within H3K27me3-enriched domains. Paradoxically, genes that typically escape XCI are hypersensitive to loss of XIST RNA and XCI erosion. This implies that XIST RNA normally restrains expression levels of these genes on the Xi. Importantly, increased X-linked gene expression upon erosion does not globally impact (hydroxy)methylation levels in hiPSCs or at imprinted regions. By exploring diverse differentiation paradigms, such as trilineage commitment and cardiac differentiation, our study reveals the persistence of abnormal XCI patterns throughout differentiation. This finding has significant implications for fundamental research, translational applications, and clinical use of stem cells. We underscore the importance of raising awareness within the stem cell community regarding XCI erosion and advocate for its inclusion in comprehensive hiPSC quality control.

## INTRODUCTION

X-chromosome inactivation (XCI) in placental female mammals ensures dosage compensation between females (XX) and males (XY) through transcriptional silencing of one X chromosome (Loda et al., 2022; Patrat et al., 2020). This process is indispensable for female survival (Penny et al., 1996; Marahrens et al., 1997), being established in post-implantation embryos and faithfully maintained throughout life (Werner et al., 2022). The key regulator of XCI is X-inactive specific transcript (XIST), a long noncoding RNA that remains expressed only from the X chromosome randomly chosen for inactivation (reviewed in Patrat et al., 2020). XIST coats the inactive X chromosome (Xi), and recruits several RNA binding proteins (RBPs) and chromatin modifiers for stable transcriptional silencing across the entire chromosome (da Rocha & Heard, 2017; Raposo et al., 2021). Importantly, not all X-linked genes are silenced on the Xi, with ∼15-25% of them, known as escapees, evading inactivation (Carrel & Willard, 2005; Tukiainen et al., 2017; Werner et al., 2022). *XIST* continues to be expressed in all somatic cells where it plays a role in the maintenance of XCI (Richart et al., 2022; Yu et al., 2021). However, in clinically-relevant female human pluripotent stem cells (hPSCs), namely embryonic stem cells (hESCs) and induced PSCs (hiPSCs) derived and cultured in conventional or “primed” conditions, *XIST* expression is recurrently lost upon cell passages (Mekhoubad et al., 2012; Vallot et al., 2015). The mechanisms underlying the silencing of *XIST* in female stem cell cultures remain poorly understood.

Loss of *XIST* expression leads to an irreversible and progressive re-activation of the Xi, a process commonly known as XCI erosion (Mekhoubad et al., 2012; Nazor et al., 2012; Sahakyan et al., 2018; Bansal et al., 2021). The eroded X chromosome (Xe) is partially reactivated and is characterized by the loss of H3K27me3 histone modification (Mekhoubad et al., 2012; Vallot et al., 2015) and DNA demethylation at the reactivated X-linked gene promoters (Geens et al., 2016; Patel et al., 2017; Fukuda et al., 2021; Bansal et al., 2021). Cell passage is the strongest risk factor for XCI erosion (Geens et al., 2016; Patel et al., 2017; Silva, Pereira, Raposo, et al., 2021; Fukuda et al., 2021), with the speed of erosion being influenced by the culture conditions (Cloutier et al., 2022). Measures to either prevent or correct XCI erosion are necessary to maintain female hPSCs with a correct X-linked gene dosage. Indeed, XCI erosion can be minimized using xenofree and feeder-dependent classical hESC medium (Cloutier et al., 2022), but prolonged culture under these conditions may still make cells vulnerable to XCI erosion. Targeting *XIST* promoter region by CRISPR-Cas9 gene editing can restore *XIST* expression and overcome erosion, however, this depends on the error-prone mechanism of Non-Homologous End Joining (NHEJ) (Motosugi et al., 2022). XCI erosion can also be reversed by resetting hPSCs to the naïve pre-XCI state, followed by a transition back to the primed state, but maintenance of XCI is limited to a certain number of passages (Agostinho de Sousa et al., 2023). Therefore, no current methods ensure permanent prevention or correction of XCI erosion.

X-linked gene activity of the Xe typically falls between the levels observed for the Xa and the Xi (Bansal et al., 2021). The presence of Xas in naive hPSCs (XaXa) is accompanied by a global decrease in DNA methylation levels which erases methylation marks at imprinted regions (Klobučar et al., 2020; Theunissen et al., 2016). Moreover, global DNA demethylation has been linked to advanced stages of XCI erosion in female hiPSCs (Bansal et al., 2021). Given the link between X-linked gene dosage and global DNA methylation, it is important to define how the degree of erosion impacts phenomena dependent on DNA methylation. This is particularly important for imprinted loci, which are often irreversibly dysregulated in hPSCs (Bar & Benvenisty, 2019; Nazor et al., 2012), with potential long-term consequences for functionality and fitness of hPSC-cell derivatives.

A key concern for safe hPSC use in clinical applications is how XCI erosion affects their differentiation properties. Previous research indicates differences in differentiation ability and cell fate decisions between eroded and non-eroded hiPSCs. For instance, hiPSCs with no *XIST* expression exhibit more immature differentiation in teratoma assays, suggesting a significant role for XCI erosion in differentiation ability (Anguera et al., 2012). Moreover, XCI erosion in hiPSCs influences neuronal potential and cell fate decisions during cardiac differentiation (D’Antonio-Chronowska et al., 2019; Motosugi et al., 2022). XCI erosion also impacts the use of female hiPSCs with X-linked mutations for disease modeling by inadvertently activating the non-mutated allele on the Xi (Mekhoubad et al., 2012). Another important question is whether erosion across the Xi itself changes during differentiation. While eroded hPSCs typically fail to regain *XIST* expression upon differentiation, the extent to which XCI erosion is maintained, rescued, or amplified across the Xe remains unclear due to inconsistent findings across previous studies (Cloutier et al., 2022; Mekhoubad et al., 2012; Motosugi et al., 2022; Patel et al., 2017; Vallot et al., 2015).

In this study, we explored the dynamics of erosion in a collection of female hiPSCs, which includes isogenic pairs expressing high levels of *XIST* (XIST+) and low levels of *XIST* (XIST-). We employed RNA FISH, RNA-sequencing (RNA-seq) and a new methodology named RNA-amplicon-seq to quantify allelic expression in these cells before and after differentiation. We unveiled that XCI erosion is frequent yet heterogeneous among hiPSCs with no impact on the global (hydroxy)methylome and on genomic imprinting. Not all X-linked genes are affected by erosion and specific traits augment the likelihood of a gene to erode. Unexpectedly, genes that already escape XCI normally are hypersensitive to erosion. Importantly, the variability of XCI status among hiPSCs persists through differentiation. Our findings emphasize the importance of drawing attention to XCI erosion within the stem cell community and advocate its inclusion in hiPSC quality control given their implications for their basic, translational, and clinical applications.

## RESULTS

### Faulty XCI is frequent in female hiPSCs

Erosion of XCI can have a major impact in the downstream application of hiPSCs. Taking advantage of our collection of female hiPSCs (Burridge et al., 2011; Chamberlain et al., 2010; Gomes et al., 2020; Klobučar et al., 2020; Silva, Pereira, Oliveira, et al., 2021; Silva, Pereira, Raposo, et al., 2021) with a wide range of cell passages, comprising 8 lines, including 2 isogenic pairs (Table S1), we set out to investigate the stability of their Xi. First, we measured *XIST* expression by RT-qPCR and showed that our female lines either express high levels (XIST+) in both ASA and ASD (at passages-P16-20) hiPSCs, intermediate levels (XIST±) in the F7 line (P37-39), or residual levels of *XIST* (XIST-) in F002 (P41-45), CD (P21-25), CE (P16-20), AG1-0 (P86-90) and iPSC6.2 (P85-87) hiPSC lines (Fig. 1A). Cell passage had an impact but was not deterministic as XCI erosion was also observed for lines with lower passages, such as CD and CE lines (Table S1). In contrast to XIST+ ASD hiPSC line, loss of *XIST* expression in CD and F002 cell lines correlated with increased methylation levels at the YY1 binding sites within exon 1 of *XIST* (Chapman et al., 2014; Fukuda et al., 2021; Makhlouf et al., 2014; Silva, Pereira, Oliveira, et al., 2021; Silva, Pereira, Raposo, et al., 2021) (Fig. S1A). To complement RT-qPCR analysis with single-cell analysis, we conducted RNA fluorescence *in situ* hybridization (RNA FISH) for XIST and XACT. XIST coats the Xi, while XACT, another non-coding RNA, coats the Xa (Vallot et al., 2013). RNA FISH was conducted in 5 cell lines: 1 XIST+ (ASD), 3 XIST-(F002, CD and CE) and 1 XIST*±* (F7). This analysis revealed that XIST is detected in 81% of cells in XIST+ ASD, 62% in XIST*±* F7 and 0% in XIST-F002, CD and CE hiPSCs (Fig. 1B), corroborating the RT-qPCR analysis. For XACT, XIST+ ASD showed monoallelic expression in 81% of cells and biallelism was never observed. In contrast, the XIST-F002, CD and CE hiPSC lines showed a high proportion of cells with biallelic expression of XACT (83%, 82% and 80% of cells, respectively). Interestingly, the XIST*±* F7 line presented cells with biallelic XACT and no XIST or monoallelic expression of both XACT and XIST (Fig. 1B).

**Fig. 1:**
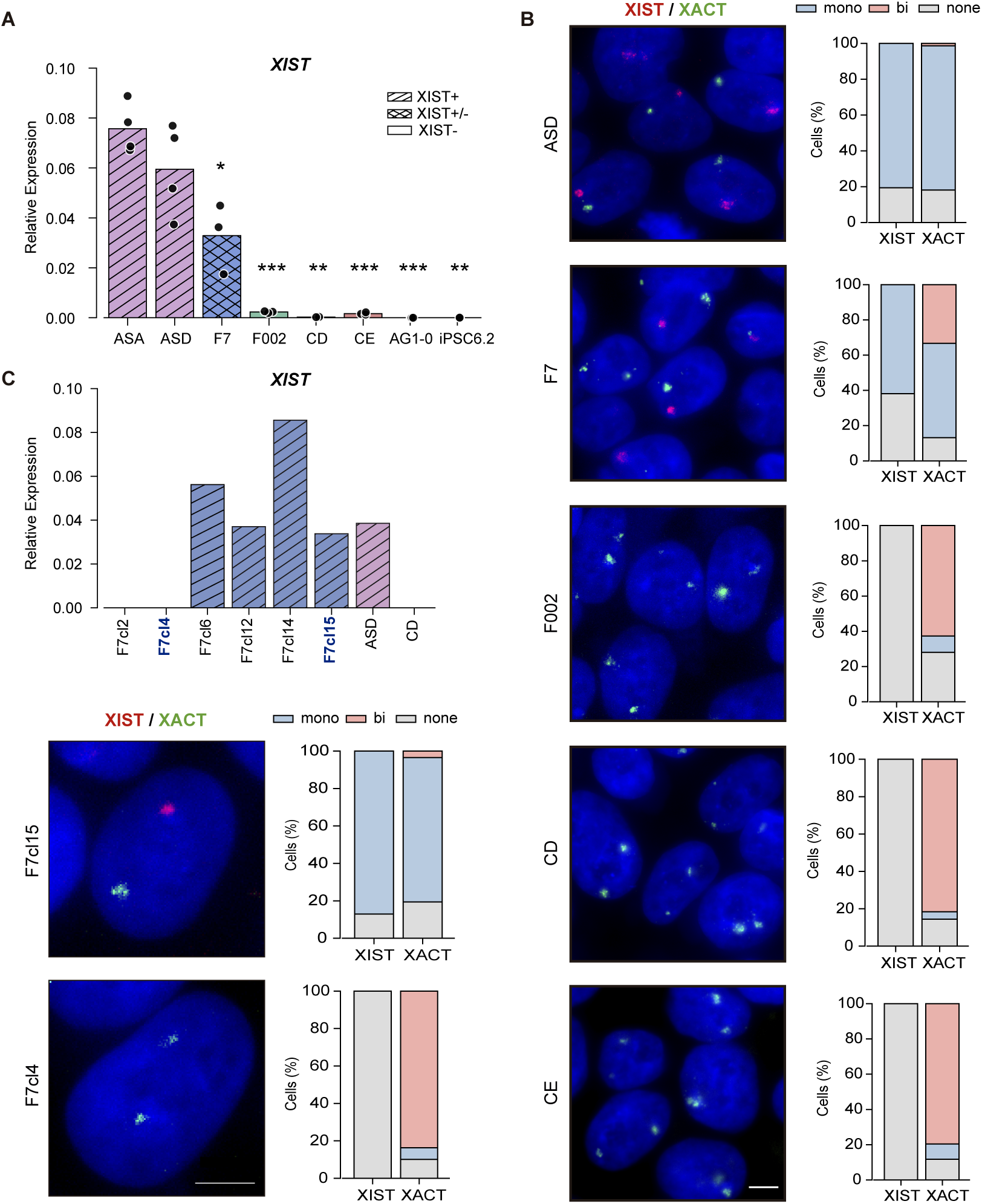
XCI status in female hiPSCs A. Barplot showing RT-qPCR analysis of *XIST* expression normalized to *GAPDH* housekeeping gene in female hiPSCs. Bars represent the average *XIST/GAPDH* relative expression. ASA, ASD and AG1-0: n=4; F7, F002 and CE: n=3; CD and iPSC6.2: n=2. Statistically significant differences between all iPSCs and ASA iPSCs are indicated as * p < 0.01; **\*\*** p < 0.001; **\*\*\*** p < 0.0001 (unpaired two-tailed Student’s t-test). **B.** Representative RNA FISH images for XIST (red) and XACT (green) in ASD, F7, F002, CD and CE hiPSCs; DNA stained in blue by DAPI; scale bar: 5 µm; graphs represent % of cells with biallelic (Bi), monoallelic (Mono), or no expression (None) of XIST and XACT; the values represent 1–4 independent experiments, where a minimum of 200 cells were counted per experiment; **C.** Above, barplot with RT-qPCR analysis of *XIST* expression normalized to GAPDH housekeeping gene in newly generated F7 cell lines (F7cl2, F7cl4, F7cl6, F7cl12, F7cl14 and F7cl15). XIST+ ASD and XIST-CD cell lines were used as positive and negative controls, respectively; n=1 for all iPSCs; Below, representative RNA FISH images for XIST (red) and XACT (green) in isogenic F7cl15 and F7cl4 hiPSCs; DNA stained in blue by DAPI; scale bar: 5 µm; graphs represent % of cells with biallelic (Bi), monoallelic (Mono), or no expression (None) of XIST and XACT; the values represent 2 independent experiments, where a minimum of 200 cells were counted per experiment. Bi-Biallelic; Mono-Monoallelic; None-No expression.

Since epigenetic states are clonally propagated, we decided to isolate XIST+ and XIST-subclones from F7 hiPSCs line by performing serial dilutions. We picked 6 viable subclones and screened them for *XIST* expression by RT-qPCR to isolate 4 F7 XIST+ subclones (F7cl6, F7cl12, F7cl14 and F7cl15) and two XIST-(F7cl2 and F7cl4) (Fig. 1C). We then selected one XIST+ (F7cl15) and one XIST-(F7cl4) subclones and analyzed the expression of XIST and XACT lncRNAs by RNA FISH. As expected, XIST was detected in 87% of cells in F7cl15 hiPSCs, while not present in F7cl4 hiPSCs (Fig. 1C). XACT was mostly monoallelic in F7cl15 (77%) and biallelic in F7cl4 hiPSCs (84%). In conclusion, these results show that our collection of hiPSC lines present different XCI states. XCI is preserved in ASD, while erosion of the Xi has occurred in F002, CD and CE hiPSCs. The F7 hiPSC line is a mosaic, containing both normal and eroded cells, from which XIST+ (*e.g.*, F7cl15) and XIST-(*e.g.*, F7cl4) subclones could be successfully isolated.

XCI erosion does not cause reactivation of X-linked genes chromosome-wide (Vallot et al., 2015; Theunissen et al., 2016; Bar & Benvenisty, 2019). Taking advantage of a panel of validated FISH probes targeting nascent transcripts, we investigated whether X-linked genes (*HUWE1*, *ATRX*, *POLA1* and *HDAC8*) are prone to reactivation upon XCI erosion. Our results show that *HUWE1*, *ATRX* and *HDAC8* genes remain monoallelic in all hiPSC lines tested, regardless of XCI status (Fig. S1B-C). Thanks to the frequent single nucleotide polymorphism (SNP) rs3088074, present in the *ATRX* gene in our hiPSCs (except for F002), we could validate clonal and monoallelic expression of this X-linked gene (Fig. S1D). Curiously, the isogenic CD and CE hiPSCs derived from the same skin biopsy (Pólvora-Brandão et al., 2018) express alternative SNPs for both genes indicating that they originated from somatic cells with opposing Xi (Fig. S1E). In contrast, *POLA1* has a different behavior: monoallelic in ASD and F7cl15 XIST+ lines (94% and 96% of cells, respectively) and consistently biallelic in F002, CD and CE XIST-lines (93%, 92% and 82%, respectively). An exception to that was the F7cl4 line for which *POLA1* remained mostly monoallelic (Fig. S1C).

Overall, our findings confirm the widespread occurrence of XCI erosion in female hiPSC cultures. Additionally, we demonstrate the resilience of certain genes (*ATRX*, *HDAC8*, and *HUWE1*) to XCI erosion, while others (*XACT*, *POLA1*) appear more susceptible. Our results also suggest potential differences in reactivation among XIST-hiPSCs.

### Different patterns of XCI erosion in female hiPSCs

To fully characterize the extent of XCI status in our female hiPSCs, we performed bulk RNA-sequencing (RNAseq) on biological triplicates of 2 XIST+ (ASD and F7cl15) and 4 XIST-(F002, F7cl4, CD and CE) hiPSCs. This includes isogenic pairs of both XIST+ and XIST-hiPSCs (F7cl15 and F7cl4) and XIST-hiPSCs with distinct eroded X chromosomes, one from maternal and the other from paternal origin (CD and CE). We first confirmed the expected pattern of *XIST* expression for each hiPSC line (Fig. 2A) and show that all hiPSC lines express high levels of pluripotency markers (Fig. S2A). We next evaluated the number of RNAseq reads mapping on the X as a proxy for erosion. A rise in RNAseq reads mapping on the X chromosome in XIST-hiPSCs was noticed when compared to XIST+ hiPSC lines. However, this increase varies substantially among different XIST-hiPSCs, ranging from 3.5% for the F002 (with residual XIST expression) to 24.6% for the CD (Fig. 2B). To better understand how this variability translates at the gene level, we conducted pairwise differential gene expression (DGE) analysis by comparing the different XIST-hiPSCs with ASD and F7cl15 XIST+ hiPSCs. As anticipated, XIST-hiPSCs exhibited a higher number of upregulated than downregulated X-linked genes, while no such trend was observed for autosomal genes when compared to both XIST+ hiPSCs (Fig. S2B-C). In line with the observed variation in read counts mapping to the X chromosome in XIST-hiPSCs, the number of upregulated X-linked genes differs among these lines, with the CD line showing the highest number of overexpressed X-linked genes (Fig. S2B-C). We also noted an elevated number of upregulated genes in ASD compared to F7cl15 (Fig. S2B-C), implying that the ASD line may exhibit initial signs of erosion, consistent with lower *XIST* levels compared to the F7cl15 line (Fig. 2A). Overall, these findings suggest that XIST-hiPSCs can exhibit quite distinct levels of XCI erosion, which is consistent with previous reports (Bansal et al., 2021; Yokobayashi et al., 2021).

**Fig. 2:**
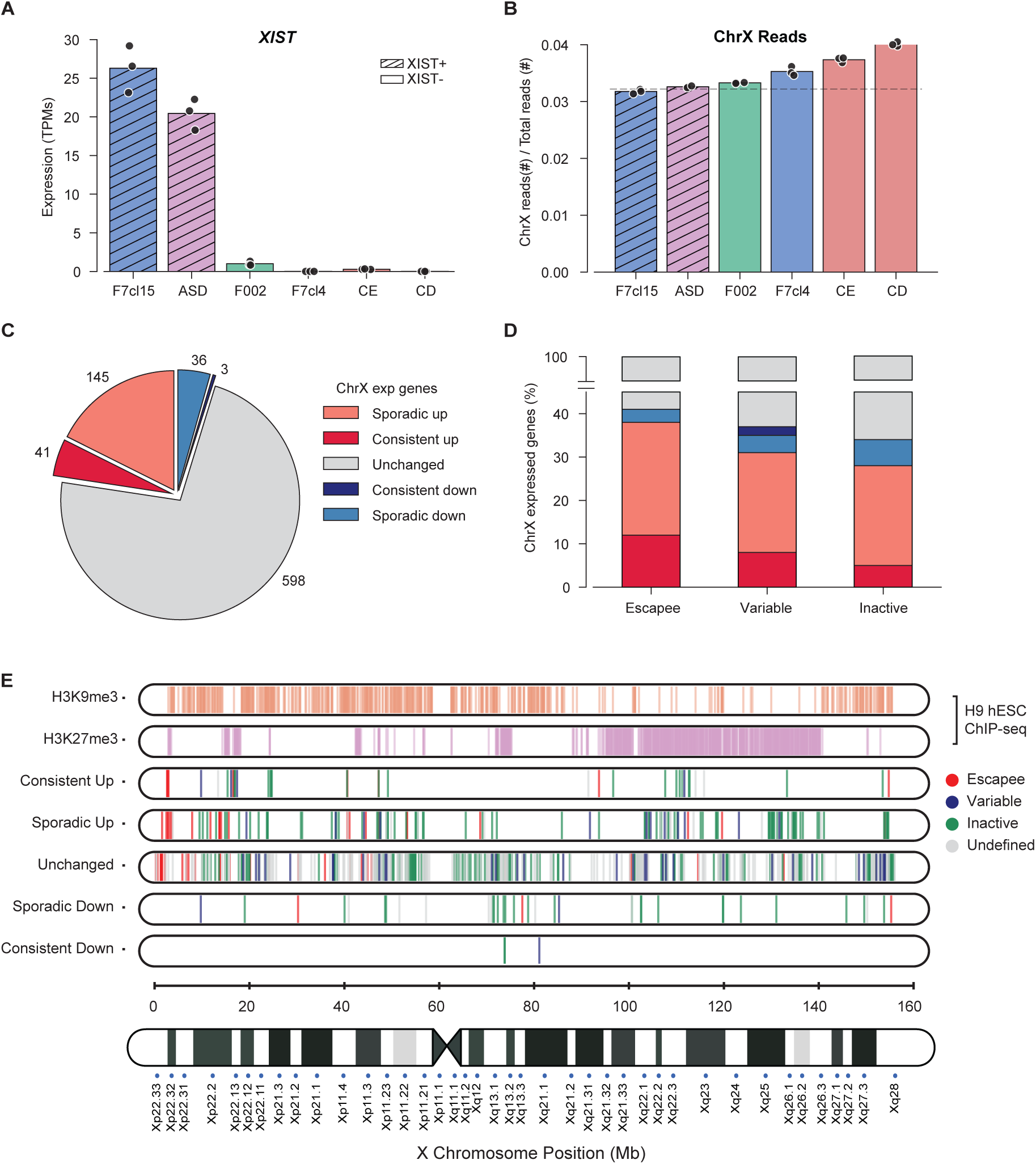
RNA-seq analysis reveals different degrees of XCI erosion in independent human iPSCs A. *XIST* expression analysis by RNAseq in F7cl15, ASD, F002, F7cl4, CE and CD hiPSCs. The graph shows the Transcripts per Million (TPMs) expression values from biological triplicates (black dots) of each sample. **B.** Barplot representing normalized X chromosome reads/total reads by RNAseq in F7cl15, ASD, F002, F7cl4, CE and CD cell lines. The graph shows the ratio between the number of chrX reads and total reads from biological triplicates (black dots) of each sample; Dashed line represents the average normalized ChrX/total reads between the two XIST+ hiPSC lines, F7cl15 and ASD. **C.** Pie chart depicting the number of consistently upregulated, sporadic upregulated, consistently downregulated, sporadic downregulated and unchanged X-linked genes in XIST-hiPSCs when compared to XIST+ hiPSCs. Consistently up/down-regulated genes: differentially expressed in 3-to-4 XIST-hiPSCs; sporadic up/down-regulated genes: differentially expressed in 1-to-2 XIST-hiPSCs; **D.** Barplot illustrating the percentage of escape, variable and inactive genes categorized according to Tukiainen 2017 and Werner 2022 (Tukiainen et al., 2017; Werner et al., 2022), within consistently and sporadic upregulated, consistently and sporadic downregulated and unchanged genes. **E.** Distribution of H3K9me3 and H3K27me3 ChIP-seq peaks in H9 hESCs (Vallot et al., 2015) and position of all genes in each category (consistently up/down, sporadic up/down and unchanged) along the X chromosome.

We then closely looked at the behavior of the different genes on the X chromosome. We observed that 41 were consistently upregulated X-linked genes (i.e., differentially upregulated in 3-to-4 XIST-hiPSCs), 145 were sporadically upregulated (i.e., differentially upregulated in 1-to-2 XIST-hiPSCs) and 598 were considered unchanged when compared to XIST+ hiPSCs (Fig. 2C; Table S2; Methods for details). Fewer genes were found consistently or sporadically downregulated (3 and 36 genes, respectively) in XIST-hiPSCs, including *XIST.* Then, we asked whether the original gene activity on the Xi (inactive, variable or escapee) prior erosion could determine the propensity of a given gene to become upregulated upon *XIST* loss. For that, we categorized X-linked genes as inactive, variable or escapee according to Tukiainen et al 2017 and reviewed by Werner et al., 2022 (Silva, Pereira, Oliveira, et al., 2021; Silva, Pereira, Raposo, et al., 2021). Among the 41 consistently upregulated genes, 34 could be categorized in these three classes (Table S2). Of those, 35% (12 genes) were identified as escapees (e.g., *GYG2*, *NAP1L3*, *TXLNG*). This percentage rises to 47% (16 genes) when variable genes are added. This is a higher proportion compared to the genome-wide number of escapees (15%) or escapees and variable genes together (∼25%) on the X chromosome. In accordance, we observed that almost 40% of the escapees expressed in hiPSCs were consistently or sporadically up-regulated, whereas the proportion decreased for variable and, even more, for inactive genes (Fig. 2D). These findings indicate that escape genes, although having evaded complete XIST-dependent silencing on the Xi, are highly susceptible to upregulation upon the loss of XIST.

We also asked whether the position along the X chromosome impacted the dynamics of X-linked gene upregulation in eroded hiPSCs. As observed in Fig. 2E, consistently upregulated genes tend to localize to specific regions of the X chromosome, notably to the short arm (especially Xp22), as well as in the central portion of the long arm (Xq22 to Xq23). We compared these regions with the genome-wide occupancy of H3K27me3 and H3K9me3 marks (Chromatin ImmunoPrecipitation-sequencing-ChIP-seq enriched regions) in XaXi H9 hESCs (Vallot et al., 2015), and found a predominant overlap between consistently upregulated genes and H3K27me3-rich domains (Fig. 2E). To corroborate this observation, we performed an enrichment analysis for each chromatin mark and found a significantly higher presence of H3K27me3 along the gene body and promoter region of all consistently upregulated genes (Fig. S2D; p-val < 0.05, Mann-Whitney U test). Sporadically upregulated genes tend to spread away from the H3K27me3-rich regions (Fig. 2E) but are still enriched for H3K27me3 (Fig. S2E), while unchanged genes are not particularly enriched for the PRC2 mark. Overall, these results confirm that X-linked genes have different sensitivities to XCI erosion with certain traits, such as escapee status, presence in Xp22 or Xq22-q23, and enrichment in H3K27me3, being predictors of erosion.

### Allele-specific profiles show increased expression following erosion of the Xi, including for escape genes

To correlate X-linked gene overexpression with reactivation of the Xe upregulated genes, we assessed allele-specific expression using RNA-seq profiles. Alleles of the two X chromosomes were discriminated using SNPs identified through whole exome-seq (WES) for the same cell lines (Methods for details). By assessing phased haplotypes of SNPs, we found around 60 expressed X-linked genes heterozygous in each cell line (Fig. 3A-B; Table S3), some of which were present across different XIST+ and XIST-hiPSC lines. For instance, the *CHRDL1* (classified as consistently upregulated gene), is considered monoallelic in both XIST+ cell lines, but becomes biallelic upon erosion (Fig. 3A). Similarly, the escapee *PRKX* (classified as sporadically upregulated gene) is monoallelic or allele-biased in XIST+ cell lines, but becomes biallelic upon erosion (Fig. 3A), showing equal expression from both alleles. On the other hand, *SLC9A7* (classified as unchanged gene), is only reactivated in the most eroded CD cell line (Fig. 3A). Importantly, the reactivation of the Xi is accompanied by an overall increased expression (bulk RNA) of the genes (Fig. 3B). The three biological replicates for each cell line, taken from three consecutive cell passages, demonstrated remarkably consistent allelic expression of the various X-linked genes, suggesting no evolution in their XCI status (Fig. 3A; Table S3). The allelic expression analysis also revealed four distinct behaviors of X-linked genes during erosion: inactive genes in XIST+ cells that become expressed by both alleles in all XIST-cells (*CHRDL1*, *SHROOM2*); genes that are only re-expressed in the CD line (*SLC9A7*, *XIAP*, *TSPAN6*, and *MAP7D3*); genes that remain monoallelically expressed in all cell lines (e.g., *CENPI*, *TRMT2B*); and escape genes exhibiting biased allelic expression in at least one of two XIST+ lines, then becoming equally expressed from both alleles in XIST-cell lines (*PRKX*, *TXLNG*, *RBPP7*, and *EIF2S3*) (Fig. 3A and Fig. S3A). Our data reveals an unequal expression pattern of escape genes from the two X chromosomes in XIST+ cells. Interestingly, this phenomenon is reversed to equal 50%:50% allelic expression in XIST-hiPSCs. These findings indicate that XIST limits the expression of escape genes on the Xi, potentially explaining why these genes show increased expression when XIST is absent in eroded cells.

**Fig. 3:**
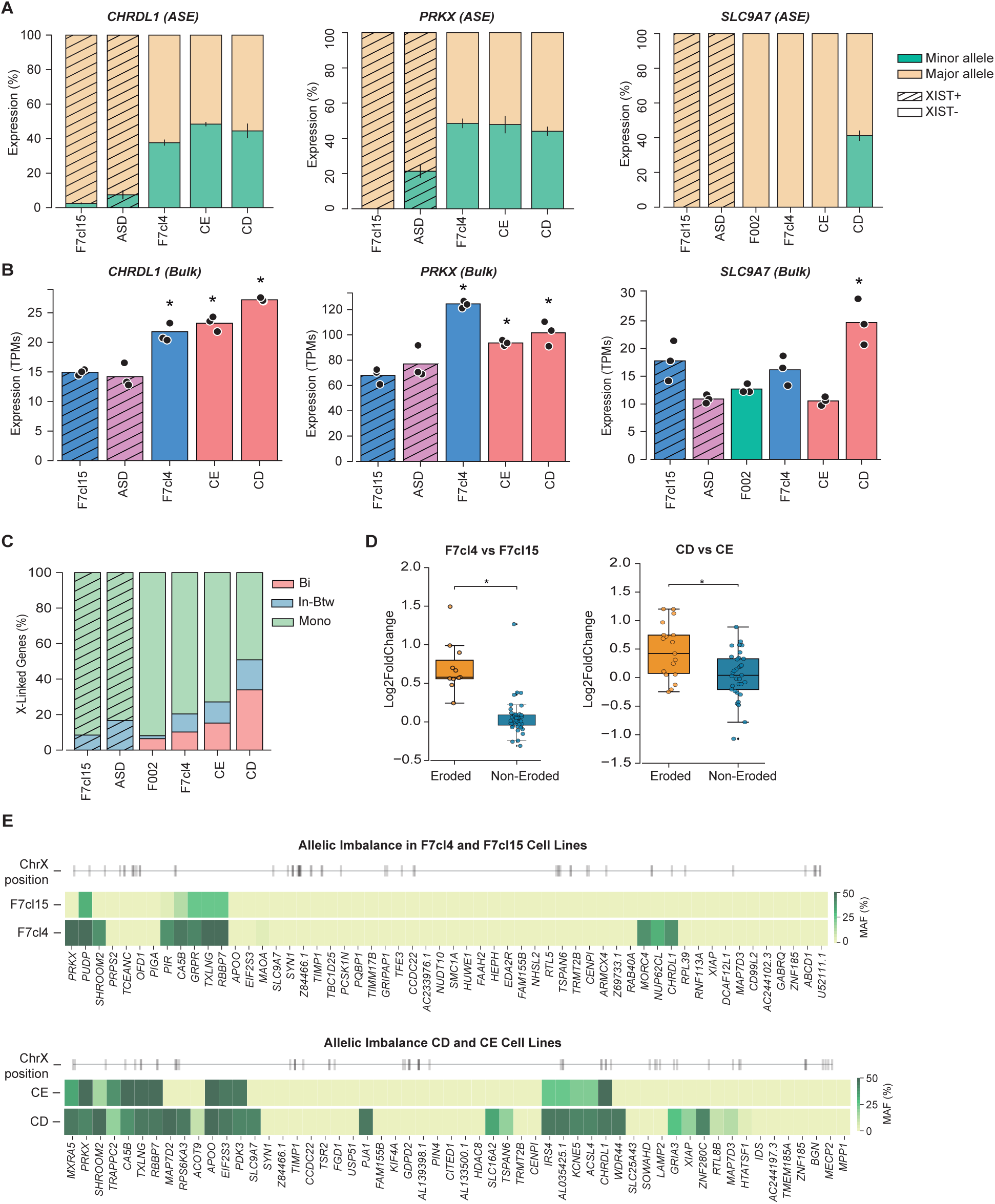
Eroded hiPSC lines show increasing number of biallelic gene expression A. Allele-specific expression (ASE) based on RNAseq analysis from representative genes containing common SNPs across different hiPSCs. Bar plots displaying the average percentage of expression ± SEM of the major and minor alleles for the X-linked genes *CHRDL1*, *PRKX* and *SLC9A7* in triplicates of XIST+ (F7cl15, ASD) and XIST-(F002, F7cl4, CE and CD) hiPSCs. **B**. Corresponding RNA bulk expression based on RNAseq analysis for these X-linked genes in the same XIST+ and XIST-hiPSCs shown in A. Bar plots represent expression in TPMs (Transcripts Per Million) for each gene in the different iPSC lines. Asterisks indicate statistical significance compared to either F7cl15 or ASD cell line (independent t-test, pval < 0.05). **C**. Percentage of biallelic (Bi), in-between (In-btw) and monoallelic (Mono) genes in XIST+ (F7cl15 and ASD) and XIST-(F002, F7cl4, CE and CE) hiPSCs. Classes were defined based on the minor allele frequency (Methods for details). **D**. Differential gene expression analysis (DGEA) between F7cl4/F7cl15 and CD/CE isogenic pairs. Genes with at least 10% increase in their minor allele frequency (MAF) were defined as eroded genes., while all the other expressed genes were considered non-eroded. Boxplots show the classification of each gene (eroded or non-eroded) against the log2FoldChange obtained from the DGEA. Asterisks indicate statistical significance (p-val < 0.01, Mann-Whitney U test). **E**. Minor allele frequency within prominent X-linked genes in isogenic hiPSC pairs, F7cl15/F7cl4 and CD/CE. Heatmap shows genes containing common SNPs in both isogenic cell lines, ordered by their position along the X-chromosome (gray lines in the ideogram).

To analyse allelic expression more globally, we divided X-linked genes in three classes based on the minor allele frequency (MAF): monoallelic (MAF<0.10), biallelic (MAF>0.40) and in-between (0.10<MAF<0.40). As expected, the XIST+ ASD and F7cl15 cell lines showed no biallelic expression and mostly monoallelic expressed genes (Fig. 3C). Consistent with the number of total X chromosome reads (Fig. 2B), we observed an increased number of in-between and biallelic genes in the eroded hiPSC lines in the following order: F002<F7cl4<CE<CD. These findings suggest that overexpression of X-linked genes in eroded hiPSCs can be attributed to increased transcriptional activity from the Xe.

Next, we took advantage of our isogenic cell line pairs (F7cl4/F7cl15 and CD/CE), each of which share the same SNPs, to conduct a more in-depth allele-specific analysis. To achieve this, we separated genes exhibiting a minimum 10% rise in expression from the preceding inactive allele (classified as eroded genes) in the most eroded line within each isogenic pair (F7cl4 and CD). This segregation was performed in contrast to the remaining genes (non-eroded genes) lacking such behavior to demonstrate that fluctuations in allelic expression from the previous Xi is correlated with gene overexpression (Fig. 3D). This is manifested, not only for the XIST-F7cl4 vs. XIST+ F7cl15 cell lines, but also for the CD vs. CE cells lines with different degrees of erosion. Not surprisingly, increased gene activity in the Xe maps to the short arm and the central portion of the long arm (Xq22 to Xq23) (Fig. 3E), as previously observed for the overexpressed genes (Fig. 2E). In summary, these findings demonstrate that the heightened expression observed in eroded hiPSC lines is primarily attributed to the increased expression from the former Xi and increases with the extent of erosion.

### No impact of XCI erosion in (hydroxy)methylation or genomic imprinting

The dosage of X-linked genes is closely linked to overall levels of DNA methylation. This association is evident in mouse ESCs or naive hPSCs with two active X chromosomes (Xas), where there is a global decrease in DNA methylation levels (Theunissen et al., 2016; Choi et al., 2017)). Hence, given that XCI erosion leads to partial reactivation of the Xi, it becomes pertinent to evaluate DNA methylation levels in eroded hiPSCs. For this analysis, we took advantage of the fact that during the course of this study, we have expanded our original cohort of hiPSCs by generating 7 new female iPSC lines from 3 different donors (Table S4) (Silva, Pereira, Oliveira, et al., 2021; Silva, Pereira, Raposo, et al., 2021). XCI status was evaluated for these cell lines by measuring *XIST* expression (Fig. S4A) complemented by XIST/XACT RNA FISH for a subset of them (Fig. S4B). We observed that only 2 out of the new 7 iPSC lines maintained *XIST* expression above the levels observed for the XIST± F7 line, while the other 5 iPSC lines were XIST-. This includes hiPSC lines of low passage number (<P15) (Fig. S4A; Table S4). Using RNA FISH, we confirmed that the XIST+ 2042cl9 hiPSC line shows a single signal for both XIST and XACT in the majority of cells, whereas the XIST-2042cl1 hiPSC line predominantly exhibit two XACT signals (Fig. S4B). This expanded our initial pool of XIST+ and XIST-female hPSC lines, now comprising 3 isogenic pairs (Table S1 and S4).

With the availability of this resource, we explored potential variations in global DNA methylation levels between XIST+ and XIST-hiPSCs. For that, we measured global 5-methylcytosine (5mC) and 5-hydroxymethylcytosine (5hmC) levels by Liquid chromatography-Mass Spectrometry (LC-MS/MS). No association between *XIST* expression and 5mC or 5hmC levels was observed in our cohort of female iPSCs (Fig. 4A-B). In line with this, our RNAseq data show no discernible differences between XIST+ and XIST-cells was found in the expression levels of the *DUSP9*, a gene previously identified as a pivotal X-linked factor influencing DNA methylation levels (Bansal et al., 2021; Choi et al., 2017), (Fig. 4C). Likewise, the presence or absence of XIST, as well as the degree of erosion, does not affect methylation patterns at imprinted regions as depicted by IMPLICON, a targeted amplicon-seq-based method to measure DNA methylation at imprinted regions (Klobučar et al., 2020; Arez et al., 2022). IMPLICON was employed across two sets of isogenic XIST+ and XIST-hiPSCs, along the isogenic pair, CE and CD, exhibiting varying degrees of erosion. Indeed, abnormal gain of DNA methylation at *DLK1-DIO3*, *PEG3* or *IGF2-H9* imprinted regions in hiPSCs, which is not seen in somatic cells (2042 and 2040 Fibroblasts-2042 and 2040 Fib), occurred independently of *XIST* expression (Fig. 4D; Table S5). In conclusion, our study demonstrates that XCI erosion within our female hiPSC cohort exerts no discernible influence on DNA (hydroxy)methylation levels or the methylation-dependent phenomenon of genomic imprinting.

**Fig. 4:**
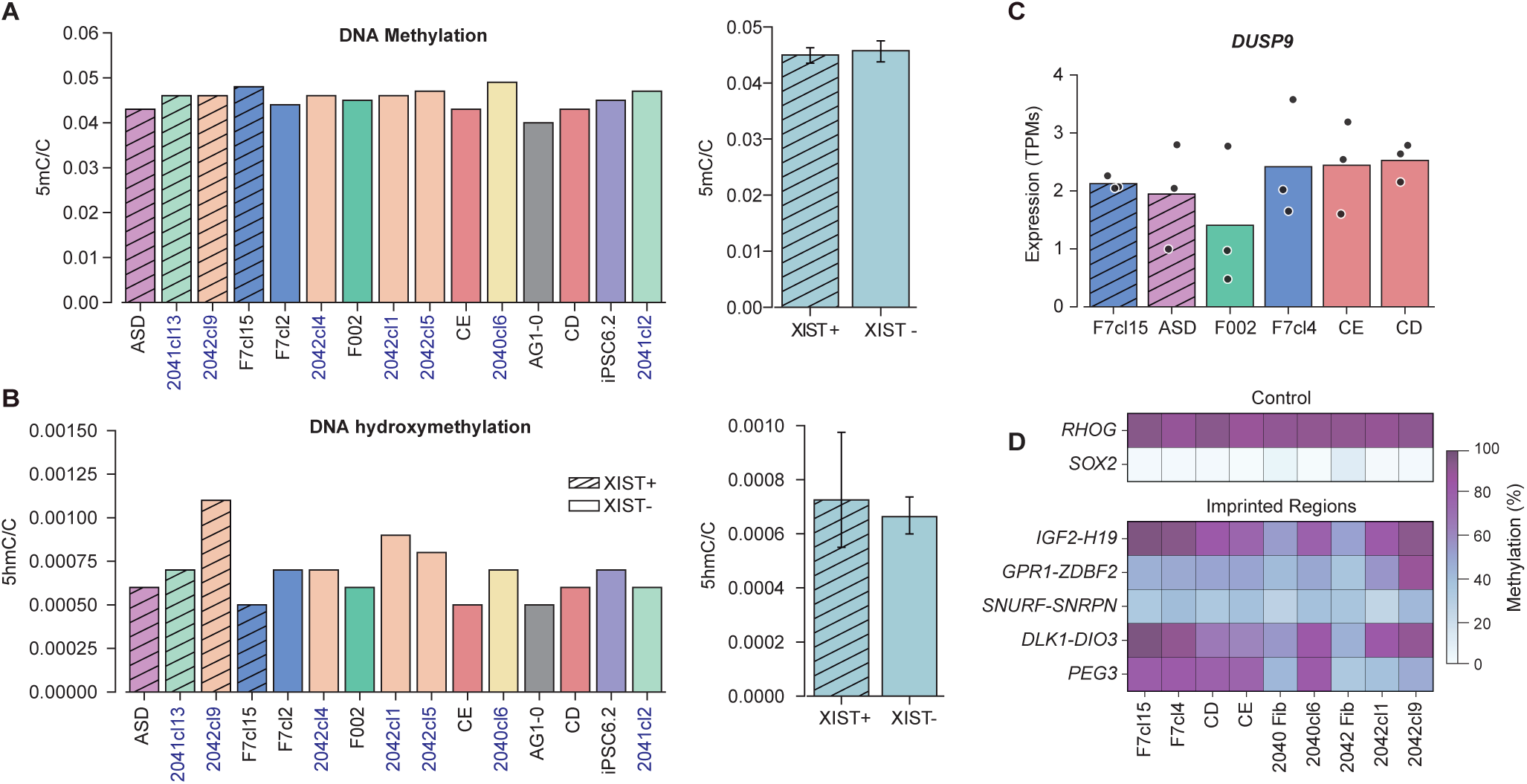
XCI Erosion Shows No Impact on Global DNA (Hydroxy)methylation and Genomic Imprinting A-B. Barplots showing global 5mC (A) and 5hmC (B) levels measured by Liquid Chromatography and mass spectrometry (LC-MS/MS). On the left, graph represents the ratio of 5mC or 5hmC per total cytosines in XIST+ (ASD, 2041cl13, 2042cl9, F7cl15) and XIST-(F7cl2, 2042cl4, F002, 2042cl1, 2042cl5, CE, 2040cl6, AG1-0, CD, iPSC6.2 and 2041cl2) hiPSCs; n=1 for all iPSCs; On the right, barplot shows the average ratio of 5mC or 5hmC ± SD per total cytosines ± SEM in XIST+ versus XIST-hiPSC lines. **C.** *DUSP9* expression data by RNAseq in F7cl15, ASD, F002, F7cl4, CE and CD hiPSCs. The graph shows the Transcripts per Million (TPMs) expression values from biological triplicates (black dots) of each sample. **D.** Heatmap representing the percentage of DNA methylation of the control loci (*RHOG* and *SOX2*) and imprinting control regions of several imprinted regions (*IGF2*-*H19*, *GPR1*-*ZDBF2*, *DLK1*-*DIO3*, *PWS*/*AS* and *PEG3*) for F7cl15, F7cl4, CD, CE, 2040cl6, 2042cl1 and 2042cl9 iPSCs as well as 2040 and 2042 fibroblasts (fib).

### Persistent XCI erosion throughout female iPSC trilineage specification

Next, we sought to explore the dynamics of Xe during differentiation. For this purpose, we first conducted trilineage commitment experiments in three isogenic pairs of XIST+ and XIST-iPSCs: F7cl15 & F7cl4, 2041cl13 & 2041cl2 and 2042cl9 & 2042cl5. These hiPSC lines were submitted to specification into ectoderm, mesoderm and endoderm for 5-to-7 days (Fig. 5A). As anticipated, all hiPSC lines demonstrated the ability to express the *PAX6* neuronal marker and downregulate *NANOG* pluripotent marker during the ectoderm differentiation (Fig. 5B). Likewise, all hiPSC lines were able to express lineage-specific markers upon differentiation to mesoderm (*BRACHYURY*) and endoderm (*SOX17*), with the exception of the XIST-2042cl5 that failed to differentiate into endoderm (Fig. S5A). It is noteworthy that *NANOG* expression remained detectable in all cell lines after endoderm specification under this short-term protocol (Fig. S5A). Overall, these data indicate that XCI erosion does not preclude trilineage specification.

**Fig.5: Erosion.**
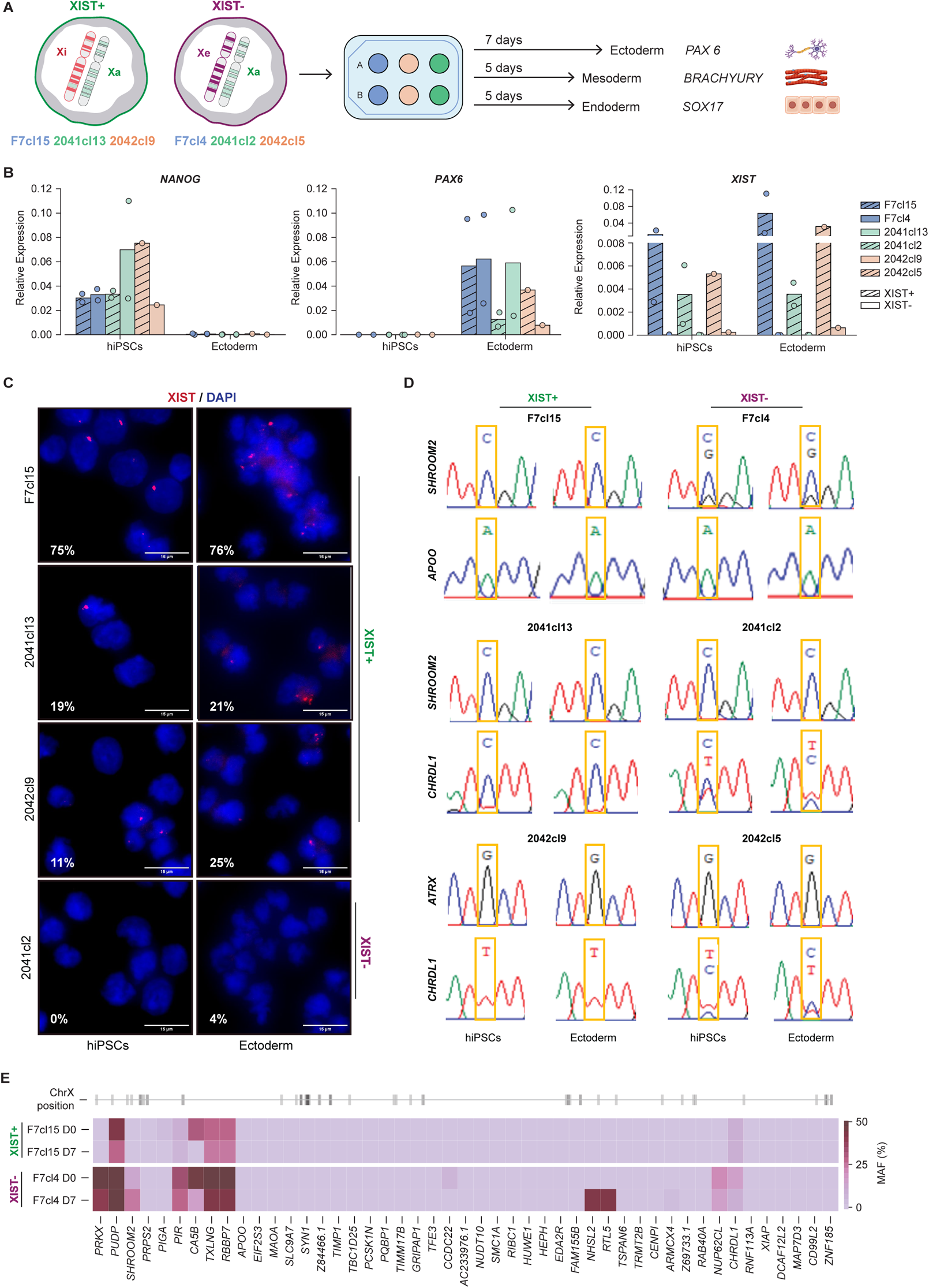
pattern persists upon ectodermal commitment. **A.** Illustration of the experimental design used for trilineage commitment of the three XIST+/XIST-isogenic hiPSC pairs (F7cl15 & F7cl4; 2041cl13 & 2041cl2; 2042cl9 & 2042cl5) into Ectoderm, Mesoderm and Endoderm. The inactive X chromosome (Xi) is marked in red, the active X chromosome (Xa) is marked in green and the eroded X chromosome (Xe) is marked in purple. **B.** RT-qPCR analysis for *NANOG* (pluripotency marker), *PAX6* (neuronal marker) and *XIST* (Xi marker) normalized to *GAPDH* housekeeping gene in F7cl15, F7cl4, 2041cl13, 2041cl2, 2042cl9 and 2042cl5 hiPSCs and after Ectoderm differentiation. Barplots represent the mean relative expression of n=2 for all samples, except 2042cl9 and 2042cl5 in both hiPSCs and Ectoderm (n=1). **C.** Representative images of XIST RNA-FISH and respective percentages of cells expressing *XIST* (red dots) in F7cl15, 2041cl13, 2042cl9 and 2041cl2 at D0 (hiPSCs) and D7 (differentiated ectodermal cells). The nuclei are counterstained with DAPI (blue). Scale bars represent 15 *μ*m. Number of cells counted: hiPSCs-F7cl15: 281, 2041cl13: 173, 2042cl9: 194, 2041cl2: 94. Ectoderm cells-F7cl15: 324, 2041cl13: 326, 2042cl9: 367, 2041cl2: 381. The values represent 1 independent experiment. **D.** Allelic expression assayed by RT-PCR followed by Sanger sequencing resourcing to informative SNPs to distinguish the two alleles. The chromatograms represent illustrative examples of the allelic expression of heterozygous X-linked genes in each XIST+/XIST-isogenic hiPSC pair at D0 (hiPSCs) and at D7 (differentiated ectodermal cells): *SHROOM2* and *APOO* gene for the F7cl15 and F7cl4, *SHROOM2* and *CHRDL1* gene for 2041cl13 and 2041cl2, and *ATRX* and *CHRDL1* gene for the 2042cl9 and 2042cl5. *CHRDL1* is not expressed in mesoderm cells. **E.** Minor allele frequency in prevalent X-linked genes between naive (D0) and differentiated (D7) cell states in F7cl15 (XIST+) and F7cl4 (XIST-) cell lines. Heatmap shows genes containing common SNPs in both states, ordered by their position along the X chromosome (gray lines in the ideogram).

We next evaluated *XIST* expression before and after trilineage commitment by RT-qPCR and RNA FISH. Reassessment of *XIST* expression after 4-6 cell passages show that XIST+ 2041cl13 and 2042cl9 hiPSC lines have undergone substantial erosion which is reflected by a reduction in the number of XIST+ cells detected by RNA FISH (19% for 2041cl13 and 11% for 2042cl9) (Fig. 5B-C). In contrast, XIST-hiPSCs maintained residual expression levels of *XIST* by both RT-qPCR and RNA FISH, while F7cl15 remained as a bona-fide XIST+ hiPSC line (75% of XIST+ cells by RNA-FISH) (Fig. 5B-C). Importantly, although the expression levels of XIST may raise slightly during ectoderm differentiation, the number of XIST expressing cells remained largely stable upon trilineage commitment and were not rescued in XIST-hiPSC lines (Fig. 5B-C; Fig. S5A-B). All in all, our results show that *XIST* expression levels do not fluctuate significantly during iPSC commitment to the three germ layers irrespective of the original levels in iPSCs.

Then, we addressed the allelic expression status of several X-linked genes in the differentiated progeny of these hiPSC lines expressing diverse levels of *XIST*. For that, we searched for eight informative common SNPs within different X-linked genes, including 6 in consistent and sporadic eroded genes (Table S6A) and qualitatively assessed allelic expression using Sanger sequencing. Overall, the *XIST*+ counterpart of each isogenic pair retains more genes with monoallelic expression (Fig. 5D; Fig. S5C; Table S6B). Despite considerable loss of *XIST*, most of the assessed genes in the 2041cl13 and 2042cl9 lines are not detected as biallelic in contrast to their XIST-counterparts. Strikingly, the original allelic expression profile found for each gene in hiPSCs, irrespective of their *XIST* expression level, is maintained upon commitment to the three germ layers (Fig. 5D; Fig. S5C; Table S6B).

To address this in more detail, we performed RNAseq for ectodermal differentiation of the XIST+/XIST-F7cl15/F7cl4 pair. As anticipated, the expression levels of pluripotent markers (*NANOG*, *POU5F1*, *PRDM14*) decreased upon differentiation, while ectoderm markers (*PAX6*, *NES*, *OTX2*) increased (Fig. S5D). Notably, *XIST* expression increased throughout differentiation in the F7cl15 line but remained absent in the F7cl4 line (Fig. S5E). We then compared the allelic expression between F7cl15 D0/D7 and F7cl4 D0/D7. With the notable exception of *NHSL2* and *RTL5* genes that became eroded upon differentiation (D7) in F7cl4 XIST-hiPSCs, the biallelically expressed genes remain the same before (D0) and after the differentiation (D7) (Fig. 5E). Additionally, the proportion of biallelic, monoallelic, and genes falling in between remained largely unchanged (Fig. S5F). An exception was the *CA5B* gene that became monoallelic in differentiated cells (Fig. 5E). As this occurs in both F7cl4 XIST-and F7cl15 XIST+, this might simply indicate that *CA5B* is a variable escape gene that evades XCI in hiPSCs but not in ectoderm cells. In conclusion, the general view is that the erosion pattern is not rescued or magnified upon ectodermal commitment, neither to endoderm or mesoderm.

### Persistent XCI erosion throughout female iPSC cardiac differentiation

Parallel to the short-term trilineage specification experiments, we conducted a long-term cardiac differentiation protocol (Lian et al., 2012) on our initial set of hiPSCs lines: ASD XIST+ and F002, CD and CE XIST-cell lines (Fig. 6A). After 15 days of differentiation, both XIST+ and XIST-cell lines were able to differentiate in contractile cardiomyocytes (Supplementary Video 1). At day 30 of differentiation, we confirmed by RT-qPCR that both XIST+ and XIST-hiPSCs downregulated the pluripotency markers, *POU5F1* and *NANOG*, and upregulated a cardiac-specific *MYBPC3* marker (Fig. 6B). As expected, *XIST* expression is maintained in ASD XIST+ hiPSCs-derived cardiomyocytes and never rescued in XIST-hiPSCs (Fig. 6B).

**Fig 6:**
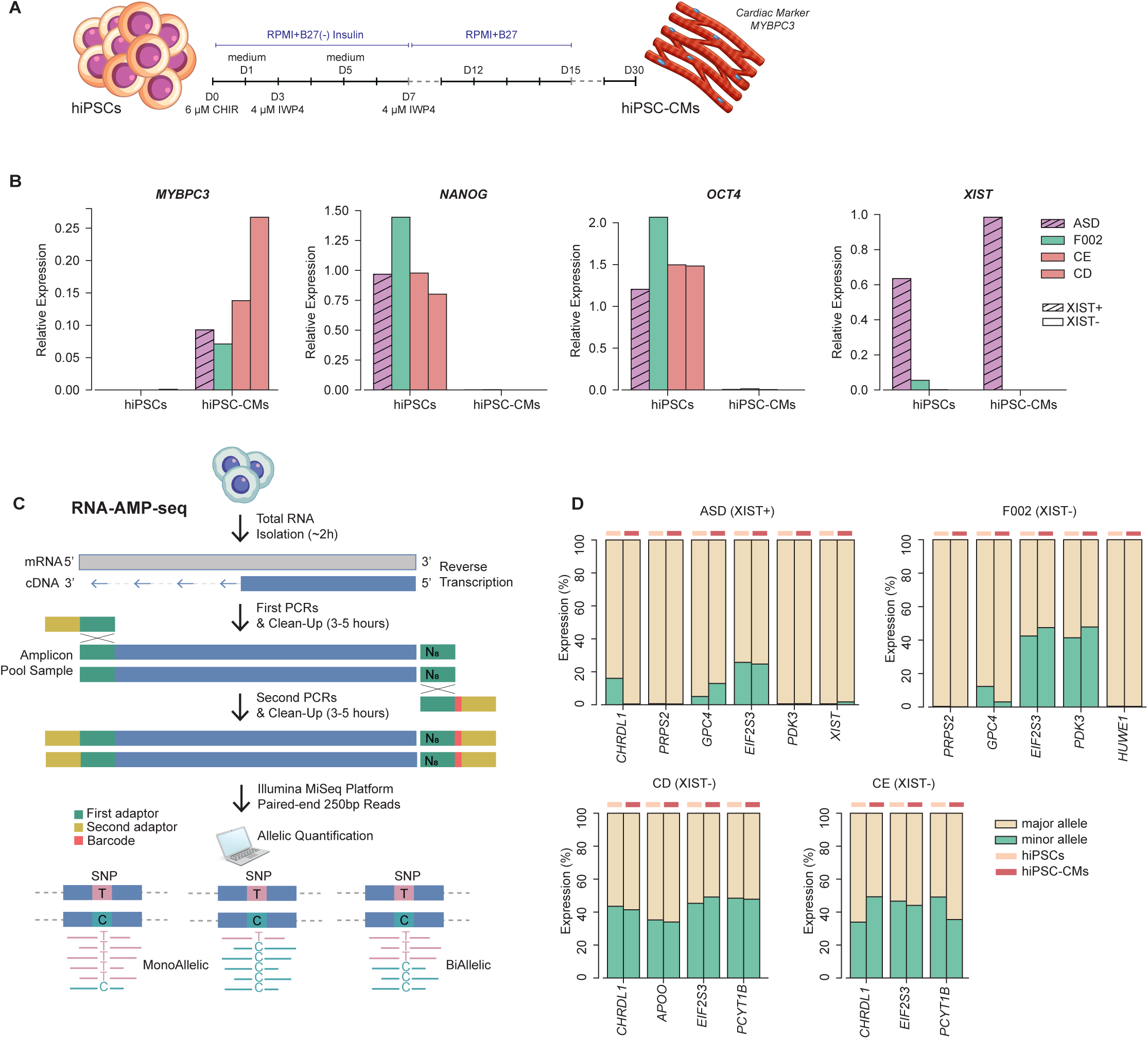
RNA-AMP-Seq reveals maintenance of allelic ratio of X-linked genes upon cardiac differentiation of female hiPSCs A. Schematic overview of the cardiac differentiation protocol (see Materials and Methods for details). **B**. Barplots with RT-qPCR analysis for *MYBPC3, NANOG, OCT4* and *XIST* expression normalized to the *U6* housekeeping gene in ASD, F002, CE and CD in hiPSCs and hiPSC-derived cardiomyocytes (hiPSC-CMs); n=1 for all the samples. **C**. Illustration of our novel RNA-AMP-Seq methodology. A first PCR amplifies each region per sample in individual reactions in the presence of adapter sequences and 8 random nucleotides (N8) for downstream deduplication of the data. After clean-up using AMPure XP magnetic beads, a second PCR completes a sequence-ready library with sample-barcodes for multiplexing. Libraries were sequenced using the Illumina MiSeq platform to generate paired-end 250 bp reads (Methods for more details) **D.** Allele-specific expression (ASE) based on RNA-AMP-seq analysis from representative genes containing common SNPs across different hiPSCs. Bar plots displaying the expression of the major and minor alleles for different X-linked genes in XIST+ (ASD) and XIST-(F002, CE and CD) hiPSCs and hIPSC-CMs.

To evaluate allele-specific expression of X-linked genes during cardiac differentiation, we devised a novel method called RNA-Amplicon-seq (RNA-AMP-Seq), drawing on our expertise with IMPLICON (Fig. 6C). This method integrates cDNA synthesis with amplicon sequencing and includes a de-duplication step to produce datasets with high coverage and precise allelic discrimination for quantifying expression of targeted genes (see Material and Methods for details). We focused on 9 X-linked genes with common SNPs in the population, including 4 consistently UP genes (*CHRDL1*, *GPC4*, *PDK3*, *PCYT1B*), 1 sporadically UP (*APOO*), 2 unchanged (*PRPS2*, *HUWE1*), 1 consistently DOWN (*XIST*) and 1 escapee (*EIF2S3*) (Table S7). First, we confirmed the added value of this method by showing the enhanced sequence coverage offered by RNA-AMP-seq by contrasting it with our RNAseq data in the same set of hiPSCs lines (Table S7). Next, we employed this method to assess the variation in allelic expression of X-linked genes with heterozygous SNPs between iPSCs and their cardiomyocyte derivatives (hiPSC-CM). Remarkably, we observe a consistent maintenance of allelic expression ratio of X-linked genes in both XIST+ and XIST-hiPSCs during the differentiation (Fig. 6D). Similar to the findings from our previous analysis using a shorter differentiation protocol, our results indicate that XCI erosion remains stable after a longer differentiation protocol producing functionally differentiated cell types. Furthermore, this underscores the potential of expanding the RNA-AMP-Seq method to accurately investigate allele-specific expression of dozens of X-linked genes across multiple samples simultaneously.

## DISCUSSION

Our comprehensive exploration of XCI erosion in hiPSCs reveals four significant characteristics of this recurring phenomenon in female stem cell cultures: (1) XCI occurs frequently but exhibits high heterogeneity; (2) the likelihood of gene reactivation is influenced by the original gene activity in the Xi, its genomic location, and chromatin environment; (3) Partial reactivation of the Xe does not induce changes in global DNA (hydroxy)methylation levels or affect the epigenetic profile of imprinted regions; (4) The heterogeneous status of the Xe is maintained in differentiated derivatives of hiPSCs.

We observed that XCI erosion is a common occurrence in female hiPSC cultures, with varying degrees of erosion observed across different cell lines. Our iPSCs were grown in mTSER Plus medium, which might not provide protection against erosion (Cloutier et al., 2022). Nevertheless, XCI erosion is observed in any of the most commonly used human stem cell media, including Stemflex, E8, or classical hESC medium under an irradiated feeder layer (Mekhoubad et al., 2012; Vallot et al., 2015; Motosugi et al., 2022; Agostinho de Sousa et al., 2023). Therefore, our observation that XCI erosion is common in female hiPSCs aligns with previous findings performed using other medium formulations. In addition to demonstrating the high frequency of XCI erosion, our data reveals its remarkable heterogeneity. This observation aligns with recent research (Bansal et al., 2021; Yokobayashi et al., 2021), yet we provide clear confirmation through allele-specific expression analysis. The variability observed underscores that XCI erosion encompasses a spectrum of states, adding complexity to this phenomenon. Although not comprehensively addressed by us in this study, the variable nature of XCI erosion might significantly impact downstream applications in both research and clinical settings. This underlines the critical need to incorporate XCI erosion assessment into routine hiPSC quality control protocols.

While acknowledging the presence of heterogeneity in XCI erosion, our investigation identifies specific traits influencing the probability of gene reactivation from the Xi. Our findings confirm that a gene’s location at the Xp22 or Xq22-q23 chromosomal regions, or its inclusion in H3K27me3-enriched domains, increased the likelihood of reactivation as noticed before (Vallot et al., 2015). A noteworthy result of our study is the increased sensitivity of escape genes to XCI erosion. While initially counterintuitive as these genes are known to escape silencing on the Xi, our allele-specific analysis shows a shift in escape gene behavior in XIST+ vs. XIST-hiPSCs. These genes, showing biased expression from one allele in XIST+ cells, transitioned to equal expression from both alleles in XIST-cells. Our findings suggest that XIST normally restricts escape gene expression on the Xi when compared to the Xa. This constraint seems to be lifted when XIST is absent. Interestingly, a recent study also demonstrated preferential transcriptional upregulation of escape genes following conditional deletion of *Xist* in mouse embryonic fibroblasts (MEFs) and hematopoietic stem and progenitor cells (HSPCs) (Yang et al., 2022). Both our findings and this report support the notion that escape genes are attenuated in expression levels on the Xi. This parallels recent observations in naive human embryonic stem cells (hESCs), where XIST dampens gene expression at the chromosomal scale (Alfeghaly et al., 2023; Dror et al., 2024). A key question for future studies is whether these two phenomena of XIST-dependent attenuation of gene expression involve the same underlying molecular mechanisms.

Our study also revealed that XCI erosion does not globally impact DNA methylation and hydroxymethylation levels or alter genomic imprinting patterns in hiPSCs. This was an important issue since naive XaXa hPSCs have global hypomethylation and erasure of imprints (Arez et al., 2022; Theunissen et al., 2016). Despite the reactivation of specific X-linked genes, the overall (hydroxy)methylation profiles remained unchanged between XIST+ and XIST-hiPSC lines. In a previous study, advanced stages of XCI erosion in female hiPSCs was associated with global DNA demethylation (Bansal et al., 2021). This was linked to upregulation of the *DUSP9* gene, a key factor explaining the X chromosome impacting global methylation (Bansal et al., 2021; Choi et al., 2017). In our XIST-hiPSCs, *DUSP9* was never found eroded which may explain why no impact on global DNA methylation was perceived. Based on our findings and those of others (Yokobayashi et al., 2021), there is reason to speculate that global DNA demethylation could be a rare occurrence associated with XCI erosion, potentially induced only under specific conditions. We also explored whether XCI erosion could contribute to the methylation variation at imprinted loci in iPSCs (Bar & Benvenisty, 2019; Klobučar et al., 2020). We rule out that XCI erosion interferes with DNA methylation patterns at imprinted genes and was not a contributor for the imprinting errors also frequently observed in hiPSCs.

Previous studies showed inconsistent findings showing either a rescue (Vallot et al., 2015) or maintenance of the XCI erosion pattern upon iPSC differentiation (Motosugi et al., 2022; Patel et al., 2017). These inconsistencies may be explained by the fact that no allele-specific expression was employed in these studies or was limited to a small number of genes. Our allele-specific expression analysis on different differentiation paradigms using isogenic sets of XIST+ and XIST-hiPSCs clearly point to the maintenance of the reactivated state of eroded genes during differentiation with a few exceptions. Therefore, the heterogeneous abnormal XCI states found in female hiPSCs are kept in their differentiated derivatives with potential consequences for their cellular functionality and/or fitness.

For our allele-specific expression analysis, we used a qualitative method, Sanger sequencing, and two quantitative methods: RNAseq and a new method named RNA-AMP-seq. While RNAseq gives a comprehensive overview of the transcriptome, RNA-AMP-seq targets specific genetic variants on genes and builds datasets with allelic discrimination and higher coverage (Table S7). RNA-AMP-seq demonstrates promise as a high-throughput method for erosion screening. Compared to our current approach, which analyzes 9 X-linked genes (5 consistently/sporadically upregulated) across 8 samples, RNA-AMP-seq offers significant scalability to accommodate a much wider range of genes and samples. Additionally, the design of new primer pairs allows for straightforward customization, enabling the application of RNA-AMP-seq to other categories of monoallelically expressed genes. This includes genes subjected to imprinting or exhibiting random monoallelic expression.

In conclusion, our study advances the current understanding of XCI erosion in female hiPSCs and its implications for stem cell biology and regenerative medicine. By characterizing the frequency, and persistence of XCI erosion, we provide valuable insights that can inform the development of improved hiPSC culture and differentiation protocols. Moving forward, continued research into the molecular mechanisms and the culture conditions driving XCI erosion will be essential for optimizing the utility of hiPSCs in various biomedical applications.

## MATERIAL AND METHODS

### Ethics

hiPSC lines used in this study were either purchased or previously generated by us (Pólvora-Brandão et al., 2018; Silva, Pereira, Oliveira, et al., 2021; Silva, Pereira, Raposo, et al., 2021) (Table S1; Table S4). Written informed consents were obtained by the donor or their legal guardian and ethically approved by the Ethics committee of the Lisbon Academic Medical Center, Lisbon, Portugal (approval numbers: 535/12 and 170/18).

#### Stem Cell Culture

All the hiPSC lines used in this study were cultured with mTeSR^TM^ Plus medium (#05825, Stem Cell Technologies) in 6-well plates previously coated with Matrigel^TM^ (#354230, Corning). Medium was changed every 24/48 hours (hrs) and the cells were routinely passed using 0.5 mM EDTA (#15575020, Invitrogen) in 1x Phosphate-Buffered Saline (PBS; #21600-044, Gibco). Cells were grown at 37°C and kept in a humidified 5% CO_2_ incubator in normoxia conditions.

To freeze, hiPSCs were dissociated with 0.5 mM EDTA/1x PBS and collected with Washing medium (Dulbecco’s Modified Eagle Medium/Nutrient Mixture F-12-DMEM-F12 #11320-033, Gibco, 10% KnockOut^TM^ Serum Replacement-KSR, #10828-028, Gibco, 1% MEM Non-essential Amino Acid Solution 100x-NEAA, #11140-050, Gibco), 1 mM L-Glutamine (#25030081, Thermo Fisher Scientific), 0.1 mM β-mercaptoethanol, (#31350-010, Gibco). After 3 minutes at 1000 rotations per minute (rpm) of centrifugation, the cells pellet was resuspended with a freezing medium composed of 90% KSR and 10% of Dimethyl Sulfoxide (DMSO, #D2438, Merck). Cell vials were stored in a liquid nitrogen tank.

### RT-qPCR

Total RNA was isolated from all hiPSCs lines using NZYol^TM^ RNA Isolation Reagent (#MB18501, NZYTech) and then treated with DNaseI (#04716728001, Roche) to remove contaminating DNA and according to manufacturer’s instructions. DNaseI-treated RNA (500 ng) was reverse-transcribed using random primers and a High-Capacity cDNA Reverse Transcription Kit (#4368814, Applied Biosystems) according to the manufacturer’s instructions. Reverse Transcriptase quantitative PCR (RT-qPCR) was performed using NZYSpeedy qPCR Green Master Mix ROX (#MB22302, NZYTech) or NZYSpeedy qPCR Green Master Mix ROX Plus (#MB22202, NZYTech) in StepOne^TM^ or ViiA^TM^ 7 Real-Time PCR Systems (Applied Biosystems). All PCR reactions were done with technical duplicates or triplicates and then normalized to the *GAPDH* housekeeping gene. The primers used are listed in Table S8. The results were analyzed with the StepOne^TM^ or the QuantStudio^TM^ RT-PCR softwares. The relative expression of each gene was determined using the 2^−ΔΔCT^ method.

### PCR/RT-PCR followed by Sanger Sequencing

To verify the presence of a specific SNP in ASD, F7, F002, CD, CE, 2041cl13, 2041cl2 and 2042cl9, 2042cl5 hiPSCs, genomic DNA isolated using conventional phenol:chloroform:isoamyl alcohol (#15593-031, Invitrogen) extraction was amplified using primers in the PCR section of Table S8. To analyze relative allelic expression of X-linked genes in F7cl15, F7cl4, 2041cl13, 2041cl2, 2042cl9 and 2042cl5 hiPSCs and their ectodermal, mesodermal and endodermal derivatives, cDNA synthesized as described in RT-qPCR section was amplified by PCR using the primers in Table S8. Both PCR products (DNA or cDNA) were cleaned using the NZYGelpure kit (#MB01102, NzyTech) and sent for Sanger sequencing to STABVIDA with data visualized and analyzed using Chromas v2.6.2 software.

### RNA FISH

The templates used for probe production were the following: XIST, a plasmid containing the 10Kb exon 5-6 plasmid (Rosspopoff et al., 2023); *XACT*: RP11-35D3 Bacterial Artificial Chromosome (BAC); *ATRX*: RP11-42M11 BAC; *HUWE1*: RP11-155O24 BAC; *HDAC8*: RP11-1021B19 BAC; *POLA1*: RP11-1104L9 BAC. Plasmid or BAC probes were prepared using the Nick translation DNA labeling system 2.0 (#ENZ-GEN111-0050, Enzo) with red or green dUTPs (red: #ENZ-42844L-0050, green: #ENZ-42831L-0050, Enzo). RNA FISH was performed according to previously published protocol (Bousard et al., 2019). For probe preparation, 4 µl of probe was precipitated using 1/10 3M NaAc (#S2889, Sigma-Aldrich), sheared salmon sperm DNA (#AM9680, Invitrogen), human *COT1* DNA (#15279011, Invitrogen) and 3 volumes of ethanol (#10000652, Fisher Chemical). The pellet was resuspended in 6 µl of deionized formamide (#F9037, Sigma) and dissolved for 15 min at 37°C with agitation. Then, the probes were denatured at 75°C for 7 min and incubated at 37°C for 30 min to prevent non-specific hybridization by *COT1* DNA. The probes were co-hybridized in hybridization buffer (20% dextran sulfate (#42867-5G, Sigma), 2x saline-sodium citrate (SSC; #S6639, Sigma-Aldrich), 1 µg/µl BSA (#R396A, Promega), 10 mM vanadyl-ribonucleoside (VRC; #S1402S, New England Biolabs) overnight at 37°C. For the experiments on Fig.1B-C, Fig.S1B-C and Fig. S4B), hiPSCs were grown on matrigel-coated coverslips, while hiPSCs and ectodermal and endodermal differentiated cells in Fig. 5C and Fig. S5B were dissociated with Accutase^TM^ (#A6964, Merck) and incubated onto poly-L-lysine (#P4832, Sigma)-coated 22×22 mm coverslips (#0101050, Marienfeld) for 5 min before RNA FISH procedure. Then, cells were fixed in 3% paraformaldehyde (PFA; #043368.9M, Thermo Fisher Scientific) for 10 min at room temperature (RT) and permeabilized for 5 min in 0.5% Triton X-100 (#T9284, Sigma) with 2 mM VRC diluted in PBS on ice. Cells were then dehydrated through 3 min incubations in 70%, 80%, 95% and 100% ethanol solutions and air-dried for 10 min before hybridization with probes. Coverslips were hybridized with fluorescent-labeled probes overnight at 37°C in a humid chamber with FA/SSC solution (50% deionized formamide, 2x SSC). On the next day, washes were carried out using FA/SSC solution, three times for 7 min at 42°C, and then with 2x SSC, three times for 5 min at 42°C. Nuclei were stained with 1:10.000 dilution of DAPI 0.2 mg/mL (#D9542, Sigma-Aldrich) in 2x SSC for 3 min at RT. Coverslips were then mounted on slides with mounting media. Cells were observed with the widefield fluorescence microscope Zeiss Axio Observer (Carl Zeiss MicroImaging) using a 63x oil objective and filter sets FS43HE, FS38HE and FS49.

### Bisulfite sequencing

Genomic DNA was purified using conventional phenol:chloroform:isoamyl alcohol extraction. Bisulfite treatment was performed using the EZ DNA Methylation Gold Kit (#D5006, Zymo Research) following manufacturer’s guidelines. Bisulfite-treated DNA was amplified by PCR for the YY1 binding sites within exon 1 of *XIST* (Fig. S1A) using the primers and conditions summarized in Table S8. PCR products were cloned into the pGEM-T Easy vector (#A1360, Promega) and at least 10 clones from each sample were sequenced. Methylation analysis was performed using BiQ Analyser v2.02 (Bock et al., 2005).

### Whole Exome-sequencing (WES)

Genomic DNA from ASD, F7, F002 and CD were purified using conventional phenol:chloroform:isoamyl alcohol extraction. Genomic DNA (1.5μg) was sent to NOVOGENE that conducted whole-exome sequencing (WES). Briefly, genomic DNA was fragmented into 180–280 bp by sonication and subjected to library preparation using the Agilent SureSelect Human All Exon V6 Kit (#5190-8864, Agilent Technologies). The enriched libraries underwent paired-end 150bp sequencing on the Illumina HiSeq 2000 platform.

Raw WES data from hiPCS cell lines was preprocessed with TrimGalore v0.4.4 (Martin, 2011) to remove possible sequencing adapters and filter sequences by Phred quality scores (“-q 20 – length 75--stringency 5--trim-n--max_n 2). Reads were further aligned with bwa mem (v0.7.15-r1140) (Li, 2013) against the GRCh38 genome assembly and duplicates were flagged with GATK4 MarkDuplicatesSpark (McKenna et al., 2010). Base scores were recalibrated (GATK4 Base Quality Score Recalibration), and a germline joint calling approach was performed with GATK4 HaplotypeCaller (Poplin et al., 2018) and GenotypeGVCFs to generate raw genotype calls. Variant calling was restricted to regions covered by Agilent v6 kit. Raw variant filtering was done with GATK4 VariantFiltration by employing hard filters based on several annotations (default values following GATK recommendations). All GATK4-based analyses were executed using the GATK version 4.1.2.0. Further processing of the call set was performed with bcftools v1.9 (Li, 2011) (multiallelic sites split, indel normalization and quality filtering), where thresholds of GQ > 30, DP > 20 reads and a minimum of 7 reads of the least covered allele were required (MIN(FMT/AD > 7). Filtered variants were annotated with Ensembl VEP (McLaren et al., 2016) using the 96 release. Chromosome X variants were further selected for downstream analysis.

### RNA-sequencing (RNA-seq) library preparation and analysis

Triplicates of F7cl15, ASD, F002, F7cl4, CE and CD hiPSC lines as well as one replicate of an ectodermal differentiation series for the F7cl15 and F7cl4 hiPSC pair (F7cl4 D0 and D7 & F7cl15 D0 and D7) were used for RNA-seq. Total RNA was isolated using NYZol and then DNase I-treatment was performed to remove contaminating DNA following the manufacturer’s recommendations. RNA (1 μg) was sent to NOVOGENE where quality of the samples was verified on a 2100 Agilent Bioanalyser system. Only samples with RIN score above 9 were processed. RNA was used for 250–300 bp insert cDNA library following manufacturer’s recommendations and libraries were sequenced with NovaSeq 6000 platform using paired-end 150-bp mode.

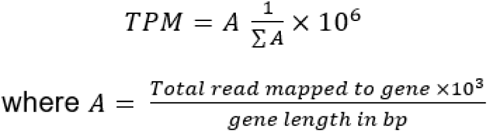

To quantify gene expression from our bulk RNAseq data (paired-ended strand-specific), we mapped the reads to the reference genome (GRCh38 assembly; release 37, GRCh38.p13) using STAR (v2.7.8a) (Dobin et al., 2013) using the-*-quantMode GeneCounts* option. The number of raw reads mapping to the X chromosome (relative to the total number of reads) was used as a proxy for erosion (Fig 2B). We calculated Transcripts per Million (TPMs) for all genes in all samples directly from the read count matrix using a custom python script.

[ueqn1]

This TPM matrix was then used to show *XIST* expression levels across samples and perform hierarchical clustering. Differential gene expression was assessed using the DESeq2 (v 1.40.2) R package (Love et al., 2014). We compared each XIST-hiPSC against each XIST+ hiPSC line and filtered differentially expressed genes (DEGs) by establishing a threshold of |log2FC| = 0.33 and an adjusted p-value < 0.05. This threshold was chosen to encompass the majority of genes that reactivate from the Xi regardless of whether they are inactive, variably expressed, or inactive genes, with an expected increase in bulk expression ranging from 1.25 to 2 times. We divided X-linked genes into five different categories according to their behavior in each comparison: consistently upregulated/downregulated (if the gene was upregulated/downregulated in three or four cell lines when compared against both ASD and F7cl15 lines); sporadically upregulated/downregulated (if the gene was upregulated/downregulated only in one or two cell lines when compared against both XIST+ controls) or unchanged (if they did not meet any of the previous criteria: e.g. gene was considered upregulated against F7cl15, but not against ASD; gene was upregulated against ASD, but downregulated against F7cl15).

Furthermore, we categorized X-linked genes according to their XCI status as inactive, variable and escapees following the classification by Tukiainen et al 2017 and then reviewed by Werner et al., 2022 (Tukiainen et al., 2017; Werner et al., 2022). We consider the classes reviewed by Werner *et al*. as reference and the classification from Tukiainen *et al*. 2017 for the remaining genes not classified by Werner *et al*., 2022.

### Allele-Specific Expression (ASE) Analysis

After conducting WES and storing gene sequence variations in a Variant Call Format (VCF), we used phASER (v.0.9.9.4) (Castel et al., 2016) for RNAseq-based phasing, enabling the generation of gene-level haplotype expression data. Only reads uniquely mapped and with a base quality ≥10 were used for phasing. For our downstream analyses, we further discarded loci with a total read depth lower than 10. This limits the number of genes in our analysis but reduces the number of false positives due to biased RNA-seq read mapping or other technical artifacts. The effect size of allelic imbalance in expression for each gene in each sample was determined using the Minor Allele Frequency (MAF), calculated as the ratio of minor allele read counts (the least common allele) to the total read counts from both alleles. We defined genes with a MAF < 0.10 as fully monoallelic, genes with a MAF > 0.40 as fully biallelic, and the remaining genes as “in-between”.

### ChIP-seq Analysis

We used ChIP-seq datasets produced by Vallot *et al*., 2015 (GEO: GSE62562) to illustrate the location of H3K27me3 (GSM1528885) and H3K9me3 (GSM1528888) marks along the X chromosome in H9 hESCs (Vallot et al., 2015). The previously processed peaks were initially converted to GRCh38 coordinates using liftOver (Kuhn et al., 2013). To further corroborate the enrichment for each mark in each category of genes (consistently and sporadic up) we counted the number of genes with at least one peak for each chromatin mark along each gene body and promoter region (gene body-4Kbp) using bedtools intersect (v2.30.0) (Quinlan & Hall, 2010). Statistical significance was assessed using the Mann-Whitney U test (Scipy package).

### 5mc/5hmc measurements by Liquid Chromatography-Mass Spectrometry (LC-MS)

Genomic DNA from hiPSCs was purified using conventional phenol:chloroform:isoamyl alcohol extraction and digested using DNA Degradase Plus (#E2020, Zymo Research) according to the manufacturer’s instructions. Nucleosides were analyzed by LC-MS/MS on a Q-Exactive mass spectrometer (Thermo Scientific) fitted with a nanoelectrospray ion-source (Proxeon). All samples and standards had a heavy isotope-labeled nucleoside mix added prior to mass spectral analysis (2′-deoxycytidine-^13^C_1_, ^15^N_2_ (#SC-214045, Santa Cruz), 5-(methyl-^2^H_3_)-2′-deoxycytidine (#SC-217100, Santa Cruz), 5-(hydroxymethyl)-2′-deoxycytidine-^2^H_3_ (#H946632, Toronto Research Chemicals). MS2 data for 5hmC, 5mC and C were acquired with both the endogenous and corresponding heavy-labeled nucleoside parent ions simultaneously selected for fragmentation using a 5 Th isolation window with a 1.5 Th offset. Parent ions were fragmented by Higher-energy Collisional Dissociation (HCD) with a relative collision energy of 10%, and a resolution setting of 70,000 for MS2 spectra. Peak areas from extracted ion chromatograms of the relevant fragment ions, relative to their corresponding heavy isotope-labeled internal standards, were quantified against a six-point serial 2-fold dilution calibration curve, with triplicate runs for all samples and standards.

### IMPLICON Library Preparation and Analysis

IMPLICON was performed as previously described (Klobučar et al., 2020) in F7cl15, F7cl4, CD, CE, 2040cl6, 2042cl1 and 2042cl9 hiPSCs, and, 2040 and 2042 fibroblasts (2040 Fib, 2042 Fib). Briefly, following bisulfite conversion, a first PCR amplifies each region per sample in individual reactions, adding adapter sequences, as well as 8 random nucleotides (N8) for subsequent data deduplication. PCR conditions and primers for this first step are listed in Table S8. After pooling amplicons for each biological sample and clean-up using AMPure XP magnetic beads (#A63880, Beckman Coulter), a second PCR completes a sequence-ready library with sample-barcodes for multiplexing. In this PCR reaction, barcoded Illumina adapters are attached to the pooled PCR samples ensuring that each sample pool receives a unique reverse barcoded adapter. Libraries were verified by running 1:30 dilutions on an Agilent bioanalyzer and then sequenced using the Illumina MiSeq platform to generate paired-end 250 bp reads using the indexing primer with the following sequence, 5’-AAGAGCGGTTCAGCAGGAATGCCGAGACCGATCTC-3’ and 10% PhIX spike-in as the libraries are of low complexity.

IMPLICON bioinformatics analysis was also performed as described (Klobučar et al., 2020), following the step-by-step guide of data processing analysis. Briefly, data was processed using standard Illumina base-calling pipelines. As the first step in the processing, the first 8 bp of Read 2 were removed and written into the readID of both reads as an in-line barcode, or Unique Molecular Identifier (UMI). This UMI was then later used during the deduplication step with “deduplicate bismark–barcode mapped_file.bam”. Raw sequence reads were then trimmed to remove both poor quality calls and adapters using Trim Galore v0.5.0 (Martin, 2011). Trimmed reads were aligned to the human reference genome in paired-end mode. Alignments were carried out with Bismark v0.20.0 and deduplication was then carried out with *deduplicate_bismark*, using the–barcode option to take UMIs into account. CpG methylation calls were extracted from the mapping output using the Bismark methylation extractor. Coverage (.cov) files were imported into Seqmonk software v1.47 (Babraham Bioinformatics; RRID: SCR_001913) for all downstream analysis. Probes were made for each CpG contained within the amplicon and quantified using the DNA methylation pipeline or total read count options.

### Trilineage Specification

Trilineage differentiation of F7cl15, F7cl4, 2041cl13, 2041cl2, 2042cl9 and 2042cl5 hiPSCs was performed using STEMdiff^TM^ Trilineage Differentiation Kit (#05230, Stem Cell Technologies) according to manufacturer’s instructions.These experiments were conducted with at least one or two replicates (Fig. 5; Fig. S5). Briefly, a density of 200,000 (mesoderm lineage) or 800,000 (endoderm and/or ectoderm lineages) of cells were plated in 12-well plates on day 0. The medium was changed daily until day 5 (mesoderm and endoderm lineages) or day 7 (ectoderm lineage). After trilineage commitment, the cells were collected with NZYol reagent to perform RNA extraction followed by RT-qPCR or RNA-seq (for F7cl15/F7cl4 ectoderm differentiation) or dissociated with accutase to perform RNA FISH and RT-qPCR experiments. The primers used are listed in Table S8.

### Cardiac Differentiation

The cardiac differentiation of ASD, F002, CD and CE hiPSCs was performed following a published protocol (Lian et al., 2012). Briefly, the hiPSCs were initially cultured in matrigel-coated plates in mTeSR1 medium (# 85850, STEMCELL Technologies) until full confluency. The differentiation was initiated by removing mTeSR1 medium and adding RPMI/B-27 without insulin (#A1895601, Thermo Fisher Scientific) and containing 6 µM of CHIR99021 (#04-0004, Stemgent), a GSK3 inhibitor. At day 3 (D3), to induce the cardiac fate of the mesendoderm progenitor cells, inhibition of canonical Wnt signaling is performed using IWP-4 (#72552, STEMCELL Technologies) a Wnt signaling inhibitor. Cardiac cells spontaneously develop into contracting cardiomyocytes when cultured in RPMI/B-27 medium and are left until day 30 of differentiation. At day 30 of differentiation (D30), the cardiomyocytes were collected with NZYol reagent to perform RNA extraction followed by RT-qPCR and RNA-AMP-seq using the primers listed in Table S8.

### RNA AMPLICON-sequencing (RNA-AMP-seq) Library Preparation and Analysis

Total RNA was isolated, DNaseI treated and reverse-transcribed as described in the RT-qPCR section of the Methods. RNA-AMP-seq was performed for the samples before (ASD, F002, CD and CE hiPSCs) and after cardiac differentiation (ASD, F002, CD, CE hiPSC-CM) using a similar procedure previously employed for IMPLICON. Briefly, a first PCR amplifies each region per sample in individual reactions, adding adapter sequences, as well as 8 random nucleotides (N8) for subsequent data deduplication. PCR conditions and primers for this first step are listed in Table S8. After pooling amplicons for each biological sample and clean-up using AMPure XP magnetic beads, a second PCR completes a sequence-ready library with sample-barcodes for multiplexing. In this PCR reaction, barcoded Illumina adapters are attached to the pooled PCR samples ensuring that each sample pool receives a unique reverse barcoded adapter. Libraries were verified by running 1:30 dilutions on an Agilent bioanalyzer and then sequenced using the Illumina MiSeq platform to generate paired-end 250 bp reads using the indexing primer with the following sequence, 5’-AAGAGCGGTTCAGCAGGAATGCCGAGACCGATCTC-3’ and 10% PhIX spike-in as the libraries are of low complexity.

RNA-AMP-seq data was first processed using standard Illumina base-calling pipelines. Briefly, the first 8 bp of Read 2 were removed and written into the readID of both reads as an in-line barcode, or Unique Molecular Identifier (UMI). This UMI was then later used during the deduplication step with “umi_tools dedup--umi-separator=’:’-I “mapped_file.bam”--paired”. Raw sequence reads were then trimmed to remove both poor quality calls and adapters using Trim Galore v0.5.0 (doi: 10.5281/zenodo.5127899, Cutadapt version 1.15, parameters:– paired). Trimmed reads were aligned to the human reference genome in paired-end mode. Alignments were carried out with STAR v2.7.11a. Deduplication was then carried out with UMI-tools v1.1 (see above). Aligned read (.bam) files were analyzed with phASER v.0.9.9.4 (Castel et al., 2016) for allelic expression quantification using the “phaser” and “phaser_gene_ae” commands.Minor Allelic Frequency (MAF) was then calculated as described above for the ASE analysis.

### Statistics and Reproducibility

The statistical methods employed in each analysis are described in their respective sections. All these statistical tests were conducted using dedicated Python packages tailored to each specific analysis.

## SUPPLEMENTARY FIGURES

### Fig. S1: Characterization of the XCI status in female hiPSCs

**A.** Schematic representation of YY1 binding sites within the *XIST* locus relative to A repeats motif. Bisulfite sequencing analysis within a region on *XIST* exon 1 containing YY1 binding sites in ASD, F002 and CD iPSCs. Each line represents the methylation profile of an independent PCR amplicon analyzed by Sanger sequencing and BiQ Analyser. White dots: unmethylated CpGs; black dots: methylated CpGs; % of methylated (Meth) CpGs was calculated as follows: average percentage of the number of methylated CpGs (black dots) / total number of CpGs (black + white dots) per cloned PCR product; the numbers in brackets represent the number of times the same amplicon was sequenced; the repetitive amplicons were only counted as one to determine the % of methylation. **B-C.** Representative RNA FISH images for *HUWE1* (red) and *ATRX* (green) in ASD, F002, CD and CE human hiPSCs (B) and for *POLA1* (red) and *HDAC8* (green) in isogenic F7cl15, ASD, F002, F7cl4, CD and CE iPSCs (C). DNA stained in blue by DAPI; scale bar: 5 µm; graphs represent % of cells with monoallelic (mono), biallelic (bi) or no expression (none). The values represent 1–2 independent experiments, where a minimum of 200 cells were counted per experiment. **D.** Summary table showing presence of heterozygosity at the SNP rs3088073 in the *ATRX* gene in F7cl15, ASD, F002, F7cl4, CE and CD human iPSCs. Bold letters refer to the presence of heterozygosity. **E.** Allelic expression of *ATRX* gene assayed by RT-PCR followed by Sanger sequencing. Chromatograms are shown for each cell line with the respective SNP highlighted in gray.

### Fig. S2: Features of X-linked genes overexpressed in eroded hiPSCs

**A.** Expression analysis by RNAseq of *PRMD14*, *NANOG* and *POU5F1* pluripotent genes in F7cl15, ASD, F002, F7cl4, CE and CD hiPSCs. The graph shows the Transcripts per Million (TPMs) expression values from biological triplicates (black dots) of each sample. **B-C.** Barplots showing the number of upregulated (red) and downregulated (blue) Differentially Expressed Genes (DEGs) in autosomes (Aut) and X chromosome (ChrX) in F7cl15, F002, F7cl4, CE and CD hiPSCs compared with ASD XIST+ line (B) and with F7cl15 XIST+ line (C). **D.** Violin plot showing enrichment analysis for H3K9me3 and H3K27me3 histone modifications in the gene body and promoter regions of consistently upregulated, sporadically upregulated and unchanged genes. Consistently and Sporadically Upregulated regions show a significant difference between both marks (pval < 0.01, Mann-Whitney U test; Cohen d ∼ 0.70).

### Fig. S3: RNAseq allele-specific expression analysis in female hiPSC lines

**A.** Allele-specific expression (ASE) based on RNAseq analysis from representative genes containing common SNPs across different hiPSCs. Bar plots displaying the average percentage of expression ± SEM of the major and minor alleles for the X-linked genes *SHROOM2*, *XIAP, TSPAN6*, *MEP7D3*, *CENPI*, *TRMT2B*, *TXLNG*, *RBPP7*, and *EIF2S3* in XIST+ (F7cl15, ASD) and XIST-(F002, F7cl4, CE and CD) hiPSCs.

### Fig S4: Analysis of XCI in the seven newly generated hiPSC lines during this study

**A.** Barplot showing RT-qPCR analysis of *XIST* expression normalized to *GAPDH* housekeeping gene in ASD, 2041cl13, 2042cl9, F7, 2042cl4, F002, 2042cl1, 2042cl5, CE, 2040cl6, AG1-0, CD, iPSC6.2 and 2041cl2 female hiPSCs. Bars represent the average *XIST/GAPDH* expression. n=1 for all iPSCs. **B.** Representative RNA FISH images for *XIST* (red) and *XACT* (green) in 2042cl9 and 2042cl1 hiPSCs; DNA stained in blue by DAPI; scale bar: 5 µm; graphs represent % of cells with monoallelic (mono), biallelic (bi) or no expression (none) of *XIST* and *XACT*; the values represent 1 experiment, where a minimum of 200 cells were counted.

### Fig S5: Erosion pattern persists upon mesodermal and endodermal commitment

**A.** Barplots showing mean relative gene expression of *NANOG* (pluripotency marker), *BRACHYURY* (mesodermal marker), *SOX17* (endodermal marker) and *XIST* (Xi marker), quantified by RT-qPCR upon normalization for *GAPDH* housekeeping gene in F7cl15, F7cl4, 2041cl13, 2041cl2, 2042cl9 and 2042cl5 hiPSCs, mesodermal and endodermal cells; n=2 for all samples, except for undifferentiated 2042 clones and differentiated (Endoderm and Mesoderm) 2041 and 2042 clones (n=1); Note that the barplots of *NANOG* and *XIST* for iPSCs are the same represented in Fig. 5B. **B.** Representative images of XIST RNA-FISH and respective percentages of cells expressing XIST (red dots) in F7cl15, 2041cl13, 2042cl9 and 2041cl2 at differentiated endodermal cells. The nuclei are counterstained with DAPI (blue). Scale bars represent 15 *μ*m. Number of cells counted: F7cl15: 327, 2041cl13: 266, 2042cl9: 243, 2041cl2: 164. The values represent 1 independent experiment. **C.** Allelic expression assayed by RT-PCR followed by Sanger sequencing resourcing to informative SNPs to distinguish the two alleles. The chromatograms represent illustrative examples of the allelic expression of heterozygous X-linked genes for each XIST+/XIST-isogenic hiPSC pair in hiPSCs and after mesoderm and endoderm specification: *SHROOM2* and *APOO* gene for the F7cl15 and F7cl4, *SHROOM2* and *CHRDL1* gene for 2041cl13 and 2041cl2, and *ATRX* and *CHRDL1* gene for the 2042cl9 and 2042cl5. Note that chromatograms for iPSCs are the same as illustrated in Fig. 5D; **D.** Heatmap showing the RNAseq expression levels for pluripotent markers (*NANOG*, *POU5F1* and *PRDM14*) and ectoderm-specific markers (*PAX6*, *NES* and *OTX2* in F7cl15 (XIST+) and in F7cl4 (XIST-) cells before (D0) and after (D7) ectodermal differentiation. **E.** *XIST* expression data by RNAseq in F7cl15 and F7cl4 before (D0) and after (D7) ectodermal differentiation (D7) (n=1). The graph shows the Transcripts per Million (TPMs) expression value. **F.** Percentage of genes classified as biallelic, in-between, or monoallelic in F7cl15 (XIST+) and F7cl4 (XIST-) cells before (D0) and after (D7) ectodermal differentiation. Classes were defined based on the minor allele frequency (Methods for details).

## LIST OF TABLES

Table S1: Description of original Cell Lines used in this study

Table S2: Differential Gene Expression Analysis (DGEA) of X-linked Genes in all cell lines Table S3: Allelic Specific Expression (ASE) of X-linked Genes in all cell lines

Table S4: Description of Cell Lines generated by reprogramming fibroblasts Table S5: Methylation analysis of 7 imprinted locus

Table S6: Informative SNPs found for the different isogenic pairs

Table S7: Allelic Specific Expression (ASE) of X-linked Genes in Cardiac Cells Table S8: Primers and Conditions

## DECLARATION OF COMPETING INTEREST

The authors declare that they have no known competing financial interests or personal relationships that could have appeared to influence the work reported in this paper.

## ACKNOWLEDGMENTS

We would like to thank the previous and current members of the S.T.d.R.’s team for helpful discussions. We also thank Felix Krueger, Maria Gouveia, Teresa Silva, Marta Furtado, and Adriana Vieira for technical help during the execution of this work. Work in S.T.d.R.’s team was supported by Fundação para a Ciência e Tecnologia (FCT) Ministério da Ciência, Tecnologia e Ensino Superior (MCTES), Portugal [IC&DT projects PTDC/BIA-MOL/29320/2017 and 2022.01532.PTDC as well as projects UIDB/04565/2020 and UIDP/04565/2020 of the Research Unit Institute from Bioengineering and Biosciences – iBB and LA/P/0140/2020 of the Associate Laboratory Institute for Health and Bioeconomy – i4HB]. A.C.R., M.A. and P.B. are supported, respectively, by SFRH/BD/137099/2018 SFRH/BD/151251/2021 and SFRH/BD/137062/2018 PhD fellowships from FCT/MCTES. P. C. is a recipient of a Marie Skłodowska-Curie Postdoctoral Fellowship (FOX-MTN-HORIZON-MSCA-2021-PF-01-01). S.T.d.R. is supported by an assistant research contract from FCT/MCTES (2021.00660.CEECIND/CP1651/CT0018).

## Notes

### Competing Interest Statement

The authors have declared no competing interest.

