## Supplemental Figures for "Erosion of X-Chromosome Inactivation in female hiPSCs is heterogeneous and persists during differentiation"

### SUPPLEMENTARY FIGURE 1

A

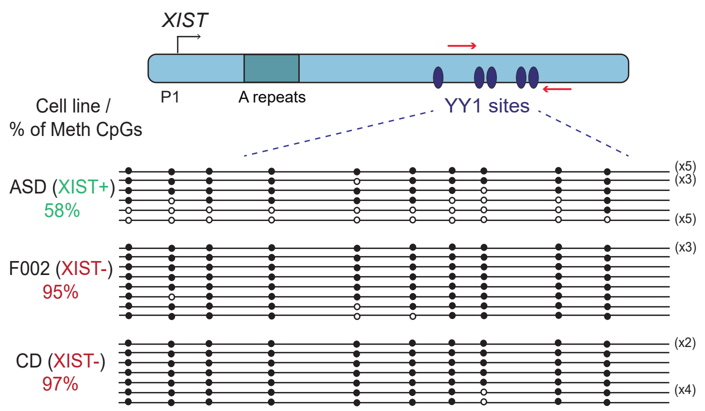


B

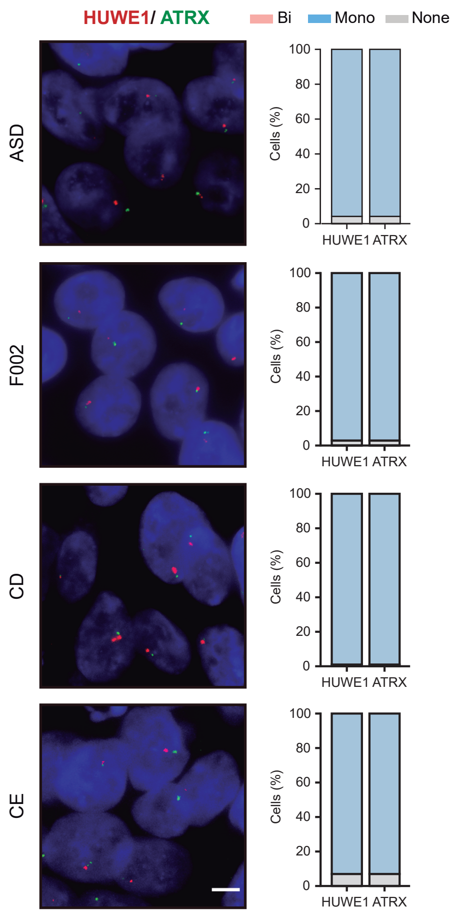


C

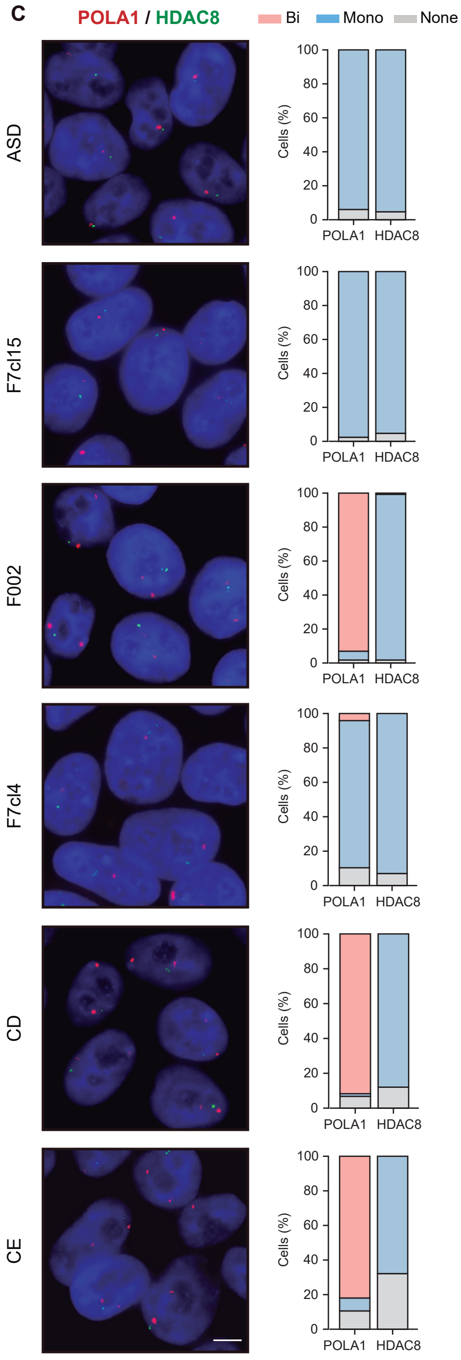


D

| Cell lines | XCI status | DNA *ATRX* (rs3088074: C/G) |
|---|---|---|
| F7cl15 | XIST+ | C/G |
| ASD | XIST+ | C/G |
| F002 | XIST- | C/C |
| F7cl4 | XIST- | C/G |
| CE | XIST- | C/G |
| CD | XIST- | C/G |

E

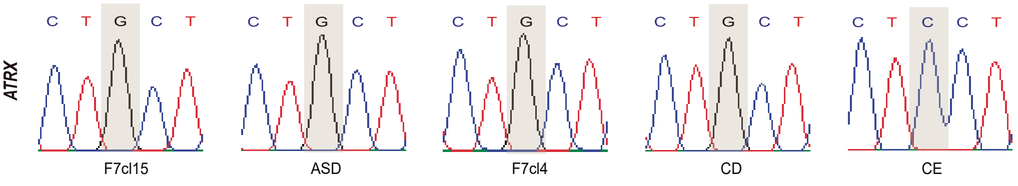

### SUPPLEMENTARY FIGURE 2

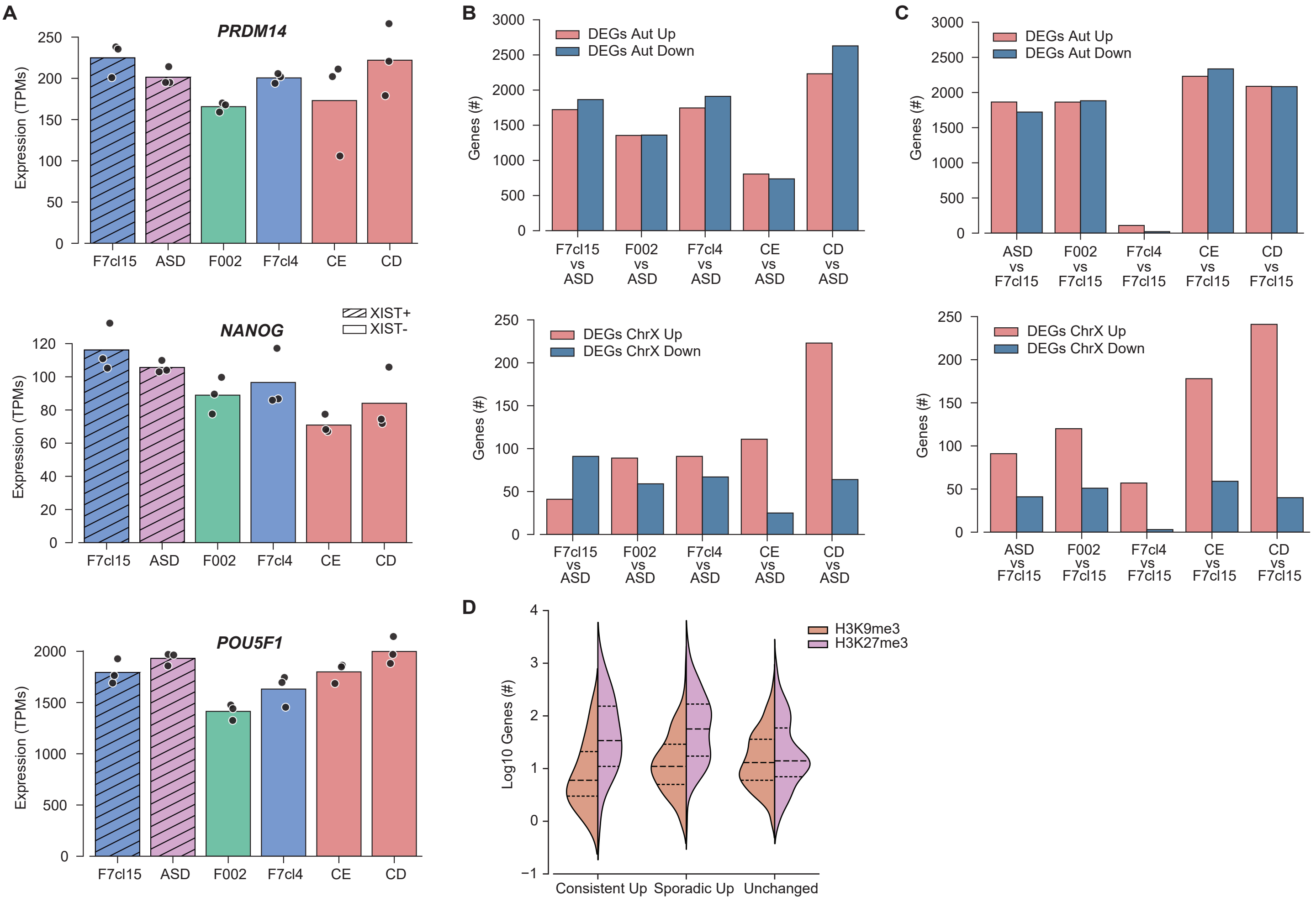

### SUPPLEMENTARY FIGURE 3

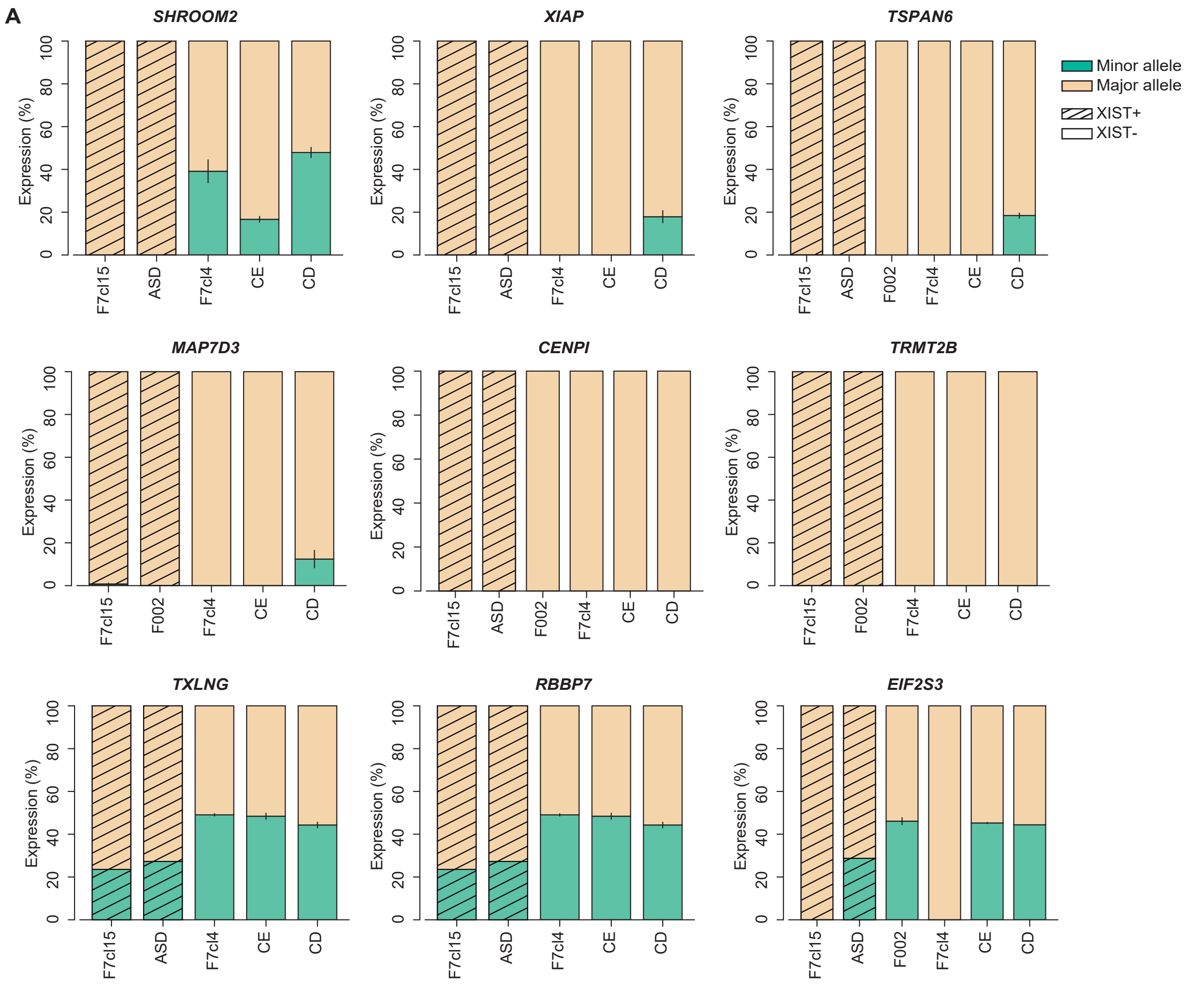

### SUPPLEMENTARY FIGURE 4

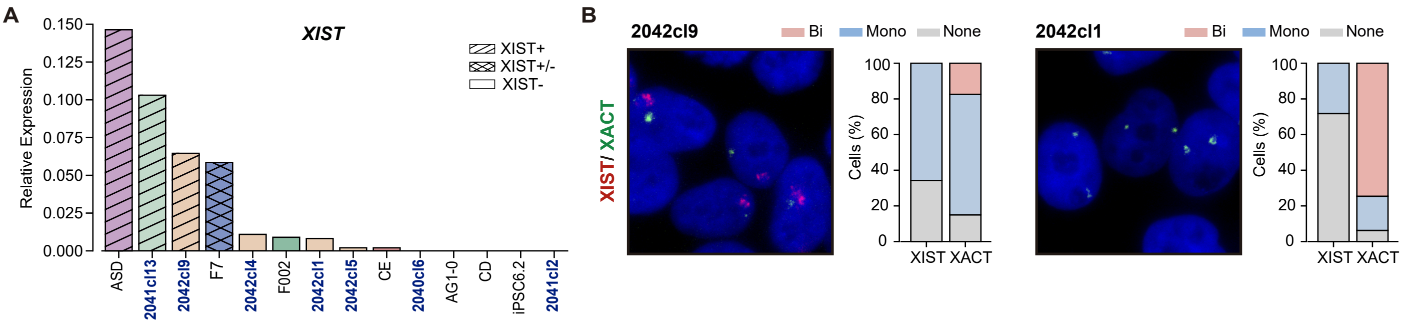

### SUPPLEMENTARY FIGURE 5

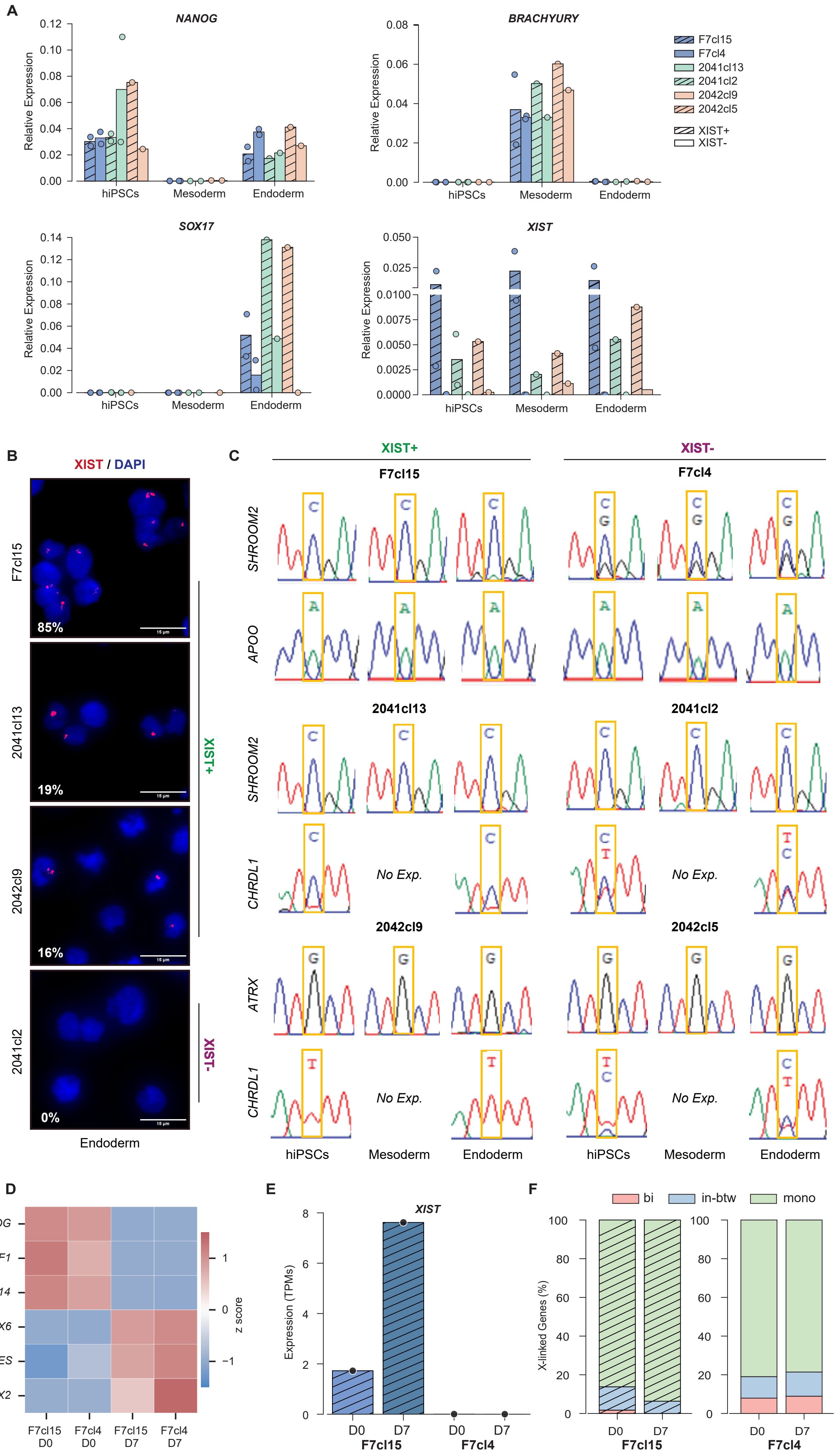
